# Normal Breast Tissue (NBT)-Classifiers: Advancing Compartment Classification in Normal Breast Histology

**DOI:** 10.1101/2025.04.27.649481

**Authors:** Siyuan Chen, Mario Parreno-Centeno, Graham Booker, Gregory Verghese, Fathima Sumayya Mohamed, Salim Arslan, Pahini Pandya, Aasiyah Oozeer, Marcello D’Angelo, Rachel Barrow, Rachel Nelan, Marcelo Sobral-Leite, Fabio de Martino, Cathrin Brisken, Matthew J. Smalley, Esther H. Lips, Cheryl Gillett, Louise J. Jones, Christopher R.S. Banerji, Sarah E. Pinder, Anita Grigoriadis

**Author notes:** These authors contributed equally (Siyuan Chen, Mario Parreno-Centeno). **Corresponding author**: Professor Anita Grigoriadis, Cancer Bioinformatics, School of Cancer and Pharmaceutical Sciences, Faculty of Life Sciences and Medicine, King’s College London, Innovation Hub, Cancer Centre at Guy’s Hospital, London SE1 9RT, UK.

## Abstract

Cancer research emphasises early detection, yet quantitative methods for normal tissue analysis remain limited. Digitised haematoxylin and eosin (H&E)-stained slides enable computational histopathology, but artificial intelligence (AI)-based analysis of normal breast tissue (NBT) in whole slide images (WSIs) remains scarce. We curated 70 WSIs of NBTs from multiple sources and cohorts with pathologist-guided manual annotations of epithelium, stroma, and adipocytes (https://github.com/cancerbioinformatics/OASIS). We developed robust convolutional neural network (CNN)-based, patch-level classification models, named *NBT-Classifiers*, to tessellate and classify NBTs at different scales. Across three external cohorts, *NBT-Classifiers* trained on 128□x□128□µm and 256□x□256□µm patches achieved AUCs of 0.98–1.00. The model learned independent normal features different from those of precancerous and cancerous epithelium, which were further visualised using two explainable AI techniques. When integrated into an end-to-end preprocessing pipeline, *NBT-Classifiers* facilitate efficient downstream analysis within peri-lobular regions. *NBT-Classifiers* provide robust compartment-specific analytical tools and enhance our understanding of NBT appearances, which serve as valuable reference points for identifying premalignant changes and guiding early breast cancer prevention strategies.

## Introduction

Normal breast tissue (NBT) research is gaining momentum for the early detection of breast cancer^1^. The epithelium, organised into ducts and lobules, also referred to as terminal duct-lobular units (TDLUs), is of primary importance, as these structures are where most physiological processes and pathological alterations occur^2^. Apart from the epithelium, recent evidence suggests that the stroma, particularly the peri-lobular stroma, may also harbour precursor alterations and have a crucial role in breast cancer progression^3–6^. Therefore, in-depth histopathological research into these structures, especially in women with varying breast cancer risk, holds promise for detecting early signs of cancer initiation in breast tissues that appear “normal”. This is particularly significant for women at higher breast cancer risk, such as germline *BRCA1/2* mutation (*gBRCA1/2m*) carriers^7^.

Histopathology remains the gold standard for diagnosing cancer, relying on microscopic examination to detect tissue abnormalities. In routine practice, pathologists first survey the entire tissue section on a glass slide before focusing on pathological regions at higher magnification for more detailed assessment. Whole slide images (WSIs) enable the hierarchical storage of microscopic histological images scanned at multiple magnifications^8^. Their ease of management and ability to facilitate rapid sharing have led to their growing integration into clinical workflows^9^, as well as driving advancements in digital image analysis using platforms such as QuPath^10^. This shift has led to the recent growth of computational pathology (CPath), a rapidly evolving field that leverages artificial intelligence (AI) and deep learning, particularly convolutional neural networks (CNNs) and the latest vision transformer (ViT)-based foundation models^11^, for high-throughput analysis of large-scale WSI datasets^12–14^.

Automatically classifying tissue and decomposing individual tissue compartments into smaller pieces, namely patches, are crucial WSI pre-processing steps. Numerous CNN-based models have been developed for breast cancer^15^, including approaches that classify tissue compartments within normal tissues adjacent to tumours, such as HistoROI^16^. In contrast, few studies have specifically focused on normal breast tissues in those individuals without malignancy, with prior efforts primarily aimed at detecting and quantifying lobules^17–22^ or assessing overall tissue composition^23,24^. Nonetheless, none of these approaches has been specifically designed to provide patch-level tissue classifications on digitised WSIs of NBTs. Moreover, all these studies typically relied on a single source of NBTs, which may limit the generalisability of the resulting models^25^. Specifically, NBTs are heavily underrepresented in large annotated WSI databases. A recent literature review^26^ summarised publicly available breast haematoxylin and eosin (H&E) WSI datasets from 2015 to 2023, identifying 17 datasets comprising a total of 10,385 breast H&E WSIs. Of these, only two contain NBTs (350 female normal breast WSIs), and just one (44 WSIs) includes manual annotations. Thus, there is a clear need for a universal patch-level classification model for NBTs, developed on a curated dataset that captures real-world variability and is supported by ground-truth manual annotations.

We present CNN-based *NBT-Classifiers* to classify patches of epithelium, stroma and adipocytes at two different scales. We provide pathologist-guided manually annotated NBTs on digitised WSIs, sourced from five cohorts, including NBTs from healthy women without high-risk germline mutations who underwent reduction surgery or core biopsy, *gBRCA1/2m* carriers who underwent a risk-reducing mastectomy, as well as contralateral and ipsilateral NBTs from breast cancer patients. *NBT-Classifiers* achieved robust performance across three external cohorts by leveraging normal-specific features, which were further validated by explainable AI-based approaches^27,28^. Additionally, an end-to-end framework was integrated, outputting pseudo patch-level annotations of target regions, including lobules and their microenvironment. Collectively, *NBT-Classifiers* provide a primary tool for studying different tissue compartments at sub-tissue level on large-scale digitised images in the context of normal breast tissue.

## Results

### Optimisation of NBT-Classifiers

In practice, pixel-level annotated digitised histological image datasets of NBTs are scarce^26^. Furthermore, the biological heterogeneity of NBTs from diverse sources is often overlooked when assembling such datasets. To address this, we collected in total 70 digitised formalin-fixed, paraffin-embedded (FFPE), H&E-stained images (n=70 patients) of NBTs from various sources, including core biopsies of healthy donors (non-*gBRCA1/2m* carriers, n=12 WSIs), women undergoing reduction mammoplasties (non-*gBRCA1/2m* carriers, n=30 WSIs), *gBRCA1/2m* carriers undergoing risk-reducing mastectomy (n=21 WSIs), and contralateral or ipsilateral normal tissues from breast cancer patients (n=7 WSIs) (**Fig. 1a**, **Supplementary Table 1**, **Methods**). These NBT WSIs were sourced from five distinct patient cohorts: the King’s Health Partners Cancer Biobank (KHP) in London (UK), the Netherlands Cancer Institute (NKI) in Amsterdam (Netherlands), the Barts Cancer Institute (BCI) in London (UK), the École Polytechnique Fédérale de Lausanne (EPFL) in Lausanne (Switzerland), and the publicly available Susan G. Komen Tissue Bank (SGK)^29^ (**Fig. 1b**). Patient age ranged from 16 to 74 years. Collectively, these digitised images capture both technical variability (differences in H&E-staining and scanning) and biological diversity (variations in tissue sources and a broad patient age range) within NBTs. Due to their greater data diversity, WSIs from the NKI and BCI cohorts were used for model training and optimisation, while WSIs from the KHP, EPFL, and SGK cohorts were held out for external validation. Expert manual annotations of epithelium, stroma, and adipocytes were performed under the supervision of a consultant pathologist (SEP) using QuPath v0.3.0 (**Fig. 1c**, **1d**, **Supplementary Fig. 1**, **Methods**). The pathology-guided annotated WSIs of NBTs are available in the nOrmal breASt tISsue Dataset (OASIS) repository, accessible at https://github.com/cancerbioinformatics/OASIS.

**Figure 1.**
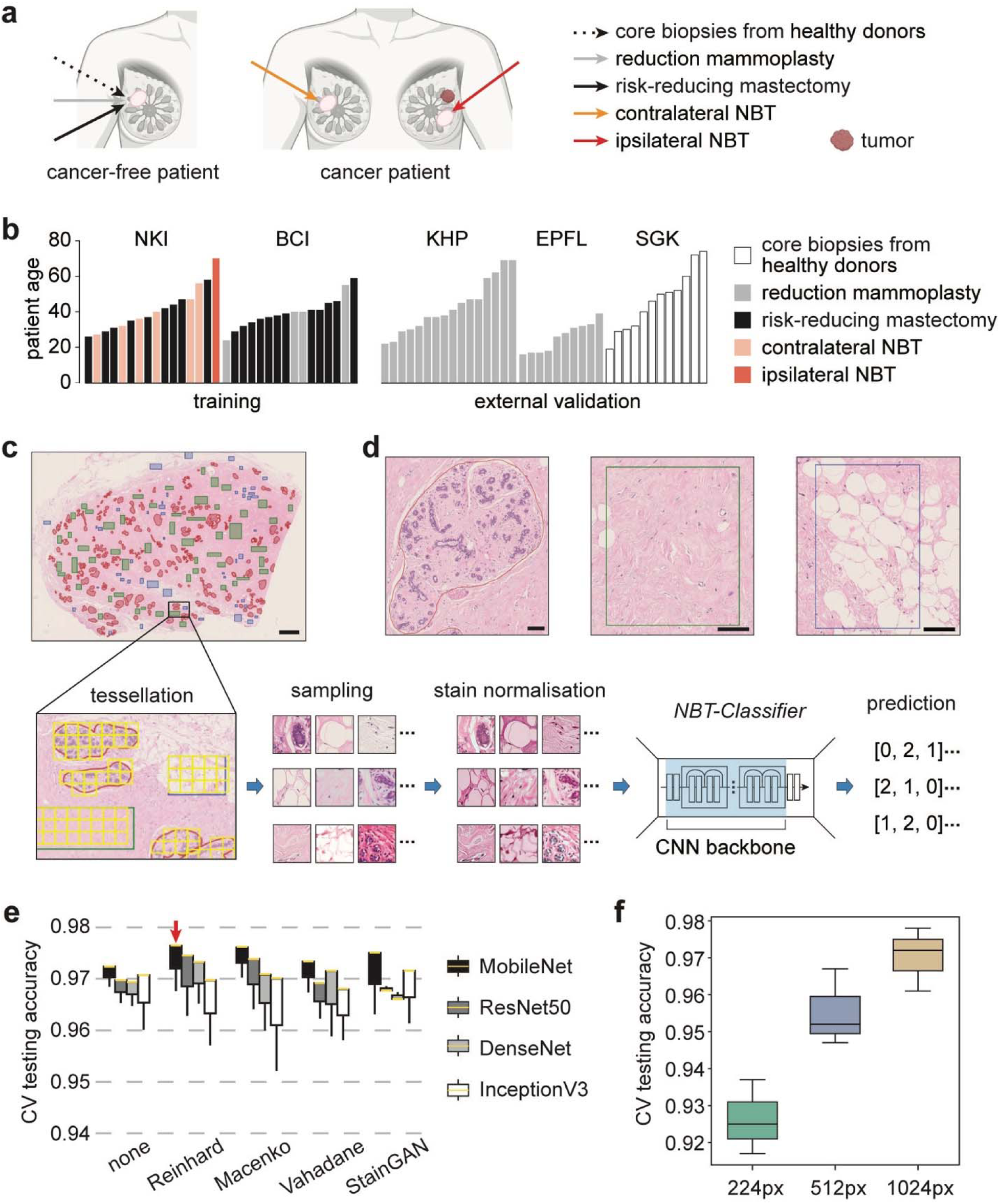
Dataset overview and optimisation of *NBT-classifiers.* **a** Schematic illustrating the sources of normal breast tissues (NBTs) used in the study. **b** Barplot depicting the distribution of whole slide images (WSIs) used for training and external validation. WSIs from the NKI and BCI cohorts were used for model training, while those from the KHP, EPFL, and SGK cohorts were reserved for external validation. Each bar represents the patient age associated with a single WSI, with colours indicating the NBT source. **c** Schematic of the manual annotation process and the training pipeline. Epithelium, stroma, and adipocytes are annotated in red, green, and blue, respectively. Scale bar, 2 mm. Annotations were tessellated into non-overlapping patches, followed by class-balanced sampling and stain normalisation for training each *NBT-Classifier*. Each model uses a CNN backbone with a classification head that predicts three tissue classes: 0 (epithelium), 1 (stroma), and 2 (adipocytes). **d** Magnified images of single annotations for epithelium, stroma, and adipocytes (from left to right). Scale bars, 100 µm. **e** Barplot of 3-fold cross-validation (CV) performance for models trained with and without different stain normalisation methods and CNN backbones. Experiments were conducted on patches tessellated at 512 × 512 pixels (0.25 µm/pixel). The yellow line in each box represents the median internal test accuracy, with whiskers extending to the minimum and maximum values. The best-performing combination is highlighted with a red arrow. **f** Barplot showing test accuracies during 3-fold cross-validation for models trained on different patch sizes. Patches were tessellated at 0.25 µm/pixel resolution and processed using the Reinhard stain normalisation method and the MobileNet backbone. The results for 224 × 224 (green, 224px), 512 × 512 (blue, 512px), and 1024 × 1024 (light brown, 1024px) pixel patches are coloured differently. The upper, middle, and lower lines of each boxplot represent the test accuracies during 3-fold cross-validation.

NBTs inherently exhibit a hierarchical organisation. Within the epithelium, structures ranging from individual epithelial cells, through to larger islands of acini, and then lobules, can be captured at fields of view (FOVs) of 64 × 64 µm, 128 × 128 µm, and 256 × 256 µm (**Supplementary Fig. 2**). Therefore, we aimed to train models enabling tissue classification at different scales to offer greater analytical flexibility for downstream analysis. To train these models, we generated patch-level datasets using QuPath v0.3.0 for the NKI and BCI cohorts. These datasets consisted of non-overlapping patches with sizes of 224 × 224, 512 × 512, and 1024 × 1024 pixels, all extracted at a resolution of 0.25 µm/pixel (**Supplementary Table 2**, **Methods**).

In the training pipeline (**Fig. 1c**), patches were subjected to stain normalisation to minimise cohort variability (**Supplementary Fig. 3**, **Methods**). The model employs a pre-trained CNN backbone for feature extraction, with a trainable classification head that predicts the probabilities of epithelium, stroma, and adipocytes (**Supplementary Fig. 4**, **Methods**). Given the critical role of stain normalisation^30^ and the selection of CNN backbone^31^, we initially performed three-fold cross-validation experiments to optimise these parameters (**Methods**). The stain normalisation methods evaluated included Reinhard^32^, Vahadane^33^, Macenko^34^, and StainGAN^35^ (**Supplementary Fig. 5**, **Methods**). For the CNN backbones, we tested MobileNet^36^, ResNet50^37^, DenseNet^38^, and InceptionV3^39^, using weights pre-trained on ImageNet^40^. These experiments were conducted using patch-level datasets containing non-overlapping 512 × 512-pixel patches from the NKI and BCI cohorts, with patches sampled in a class-balanced manner to ensure equal representation of tissue types (**Supplementary Fig. 6-7**, **Methods**). The combination of Reinhard stain normalisation and the MobileNet backbone yielded the best overall performance (**Fig. 1e**, **Supplementary Table 3**).

Configured with Reinhard stain normalisation and the MobileNet backbone, we trained models using class-balanced, non-overlapping patches of 224 × 224, 512 × 512, and 1024 × 1024 pixels from the NKI and BCI cohorts (**Supplementary Fig. 6-7**, **Methods**). The models trained on larger FOVs generally achieved higher accuracies (**Fig. 1f**, **Supplementary Table 3**), while the 224 × 224-pixel model exhibited the lowest accuracy and was more prone to misclassifying epithelium (**Supplementary Fig. 8**). Due to the suboptimal performance of the 224 × 224-pixel model, we proceeded to retrain two final versions of the *NBT-Classifier*: one using 512 × 512-pixel patches and the other using 1024 × 1024-pixel patches.

### *NBT-Classifiers* demonstrate strong generalisability underpinned by interpretable tissue representations

We then evaluated the optimised *NBT-Classifiers* on the external KHP, EPFL, and SGK cohorts using receiver operating characteristic (ROC) analysis (**Methods**). The class-specific area under the curve (AUC) values were 0.98 for epithelium, 0.98 for stroma, and 1.00 for adipocytes across all cohorts (**Fig. 2a**), with cohort-specific AUCs showing some variation (**Supplementary Fig. 9**). The 1024px-based *NBT-Classifier* achieved similarly robust class-specific AUCs of 0.99 for epithelium, 0.98 for stroma, and 1.00 for adipocytes (**Fig. 2b**), with slight variations across cohort-specific AUCs (**Supplementary Fig. 10**). For both 512px- and 1024px-based *NBT-Classifiers*, we observed no bias in the accuracy concerning patient age in external cohorts or source of the NBT (**Supplementary Fig. 11**, **Methods**).

**Figure 2.**
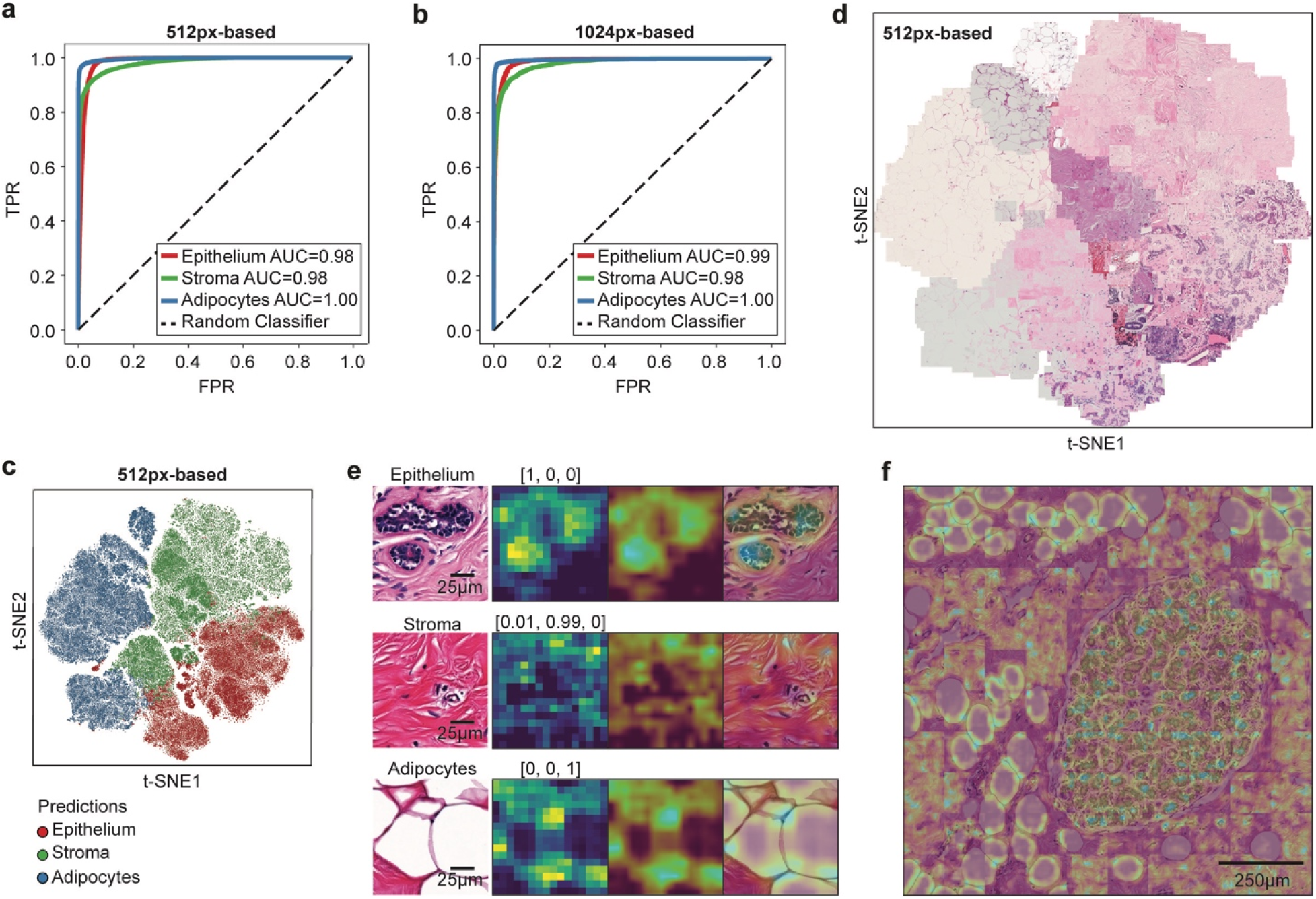
*NBT-Classifiers* demonstrate high generalisability based on robust normal breast features. **a** Tissue-specific ROC curves showing the external performance of the 512px-based *NBT-Classifier*. True positive rate (TPR) is plotted against false positive rate (FPR) at different thresholds for epithelium, stroma, and adipocytes, indicated in red, green, and blue, respectively. The corresponding area under the curve (AUC) values are presented in the legend. **b** Tissue-specific ROC curves showing the external performance of the 1024px-based *NBT-Classifier*. **c** t-SNE visualisation of the 512px-based *NBT-Classifier*’*s* predictions. Features for epithelium, stroma, and adipocytes are indicated in red, green, and blue, respectively. **d** t-SNE visualisation of H&E-stained patches of the same samples shown in panel b. For each predicted tissue class (epithelium, stroma, and adipocytes), 500 patches were randomly sampled and overlaid onto the plot based on their coordinates. **e** Class activation mapping (CAM) visualisation for the 512px-based *NBT-Classifier*. Examples of epithelium, stroma, and adipocytes patches (512 × 512 pixels, 0.25 µm/pixel) are shown in the first column. For each example, the original CAM heatmap, resized heatmap and heatmap overlaid on the original H&E patch image are shown on the right, with the predicted probabilities for epithelium, stroma, and adipocytes displayed at the top. Scale bar, 25 µm. **f** CAM heatmaps overlaid on a selected region. The region was divided into non-overlapping 512 × 512-pixel patches, for which CAM heatmaps were generated based on the predicted tissue class. The heatmaps were then resized and overlaid to highlight high-attention structures in each patch. Scale bar, 250 µm.

To dissect the underlying learned representations, we first visualised the feature representations from the last hidden layer of both models using t-distributed Stochastic Neighbour Embedding (t-SNE)^41^ (**Methods**). Patches predicted as epithelium, stroma, and adipocytes formed well-separated visual clusters (**Fig. 2c, Supplementary Fig. 12a**) that closely mirrored their respective histology (**Fig. 2d**, **Supplementary Fig. 12b**, **Methods**). We then examined the specific histological patterns driving class-specific predictions using class activation mapping (CAM)^27^ and gradient-weighted class activation mapping (Grad-CAM)^28^ (**Supplementary Fig. 13**, **Methods**). Both methods consistently highlighted biologically meaningful regions—epithelial cellular contents, collagen fibres, and adipocyte membranes—corresponding to the three tissue classes (**Fig. 2e**, **Supplementary Fig. 14a**, **14b**). When applied to a larger, representative region (5120 × 5120 pixels, 0.25 µm/pixel) encompassing all three tissue compartments, these characteristic patterns were coherently captured across non-overlapping patches, preserving spatial continuity (**Fig. 2f**). Notably, these patterns were reproducible across all cohorts (**Supplementary Fig. 15a**), including images scanned at 20x magnification from the SGK cohort (**Supplementary Fig. 15b**). When comparing the predictions with ground-truth annotations, most misclassifications occurred at the boundaries between class-specific visual clusters where histological features are less distinct (**Supplementary Fig. 12c**, **Supplementary Fig. 16a**). Correspondingly, the confidence of model’s predictions, approximated by the probability associated with the predicted tissue class, was notably lower at these boundaries (**Supplementary Fig. 16b**). We then applied CAM and Grad-CAM to assess patches with low-confidence predictions. The resulting heatmaps for all three tissue classes showed that the model’s attention was proportionally distributed across mixed tissue regions (**Supplementary Fig. 17**). This suggests that the misclassifications are not due to a bias in the model’s innate judgement, but rather to ambiguity in the testing patches where tissue classes are less clearly defined. Together, these analyses demonstrate the strong generalisability of the *NBT-Classifiers*, underpinned by interpretable and biologically grounded representations of normal breast tissue architecture.

### Training exclusively on normal tissue enables learning of distinctive features in the normal breast

Because there are no CNN models trained exclusively on normal breast tissue, we benchmarked the 1024px-based *NBT-Classifier* against HistoROI^16^, a ResNet18-based^37^ model trained on mixed breast tissue WSIs, ranging from histologically normal, noncancerous, precancerous, to cancerous tissues^42,43^, to classify epithelium, stroma, adipocytes, along with lymphocytes, miscellaneous tissues, and artefacts (**Methods**). HistoROI demonstrated comparable AUCs for epithelium (0.99 vs 0.99), while performance was lower for stroma (0.97 vs 0.98) and adipocytes (0.95 vs 1.00) (**Fig. 3a**, DeLong’s test, P<0.0001, **Methods**). This performance gap was more pronounced in accuracy metrics: HistoROI achieved 80%, 91%, and 72% for epithelium, stroma, and adipocytes, respectively, lower than the 87%, 94%, and 96% attained by the *NBT-Classifier* (**Supplementary Fig. 18**).

**Figure 3.**
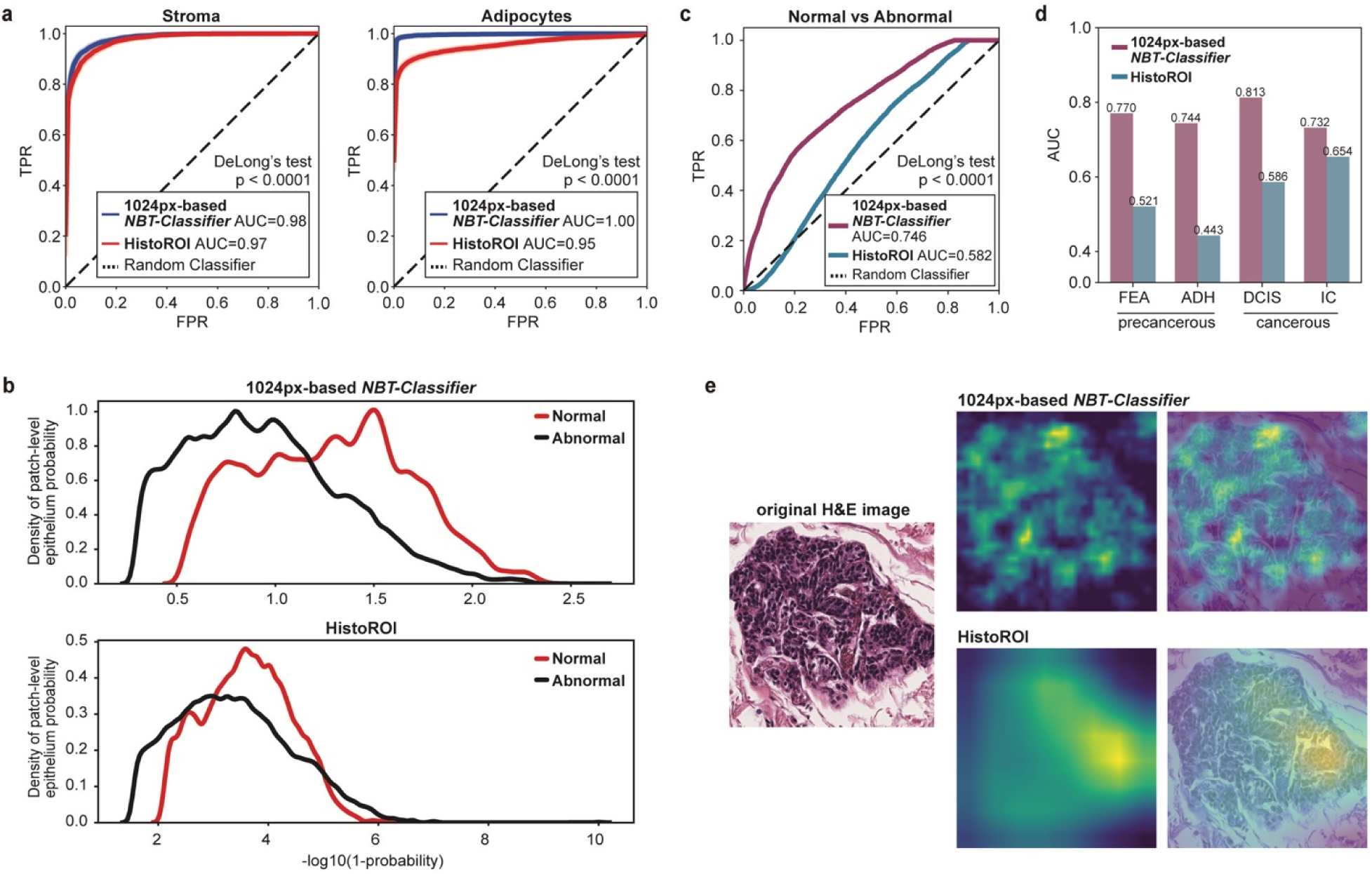
*NBT-Classifiers* capture features unique to normal breast epithelium. **a** Tissue-specific ROC curves comparing the external performance of the 1024px-based *NBT-Classifier* and HistoROI in classifying stroma and adipocytes. AUC values for each model and tissue type are listed in the legend. Differences in AUC were evaluated using DeLong’s test. **b** Distribution of epithelial probability scores for normal versus abnormal breast tissues. Panels show probability density estimates for the 1024px-based *NBT-Classifier* (top) and HistoROI (bottom). Abnormal□=□ADH, DCIS, FEA, IC, PB, and UDH lesions. **c** ROC curves for normal versus abnormal tissue classification using *NBT-Classifier* and HistoROI. Corresponding AUC values are indicated in the figure. DeLong’s test, p < 0.0001. **d** Barplots showing AUC values for normal tissue versus precancerous (FEA, ADH) and cancerous lesions (DCIS, IC) using *NBT-Classifier* and HistoROI. **e** Grad-CAM visualisation of regions of high attention contributing to epithelium predictions. For a representative epithelium patch (1024 × 1024 pixels, extracted at 40× magnification), Grad-CAM was applied to both the 1024px-based *NBT-Classifier* and HistoROI. For each model, the Grad-CAM heatmap and corresponding H&E overlay are shown.

The fundamental difference between the 1024px-based *NBT-Classifier* and HistoROI lies in their training datasets: HistoROI was developed using mixed breast tissues, while *NBT-Classifiers* were trained exclusively on normal breast tissue. This raised the question: could *NBT-Classifiers* capture unique architectural features of normal breast tissue, distinguishing it from abnormal tissues? To test this hypothesis, we curated a patch dataset using the same dataset employed to develop the HistoROI model^16,42,43^ (**Methods**). In total, 56,000 1024 × 1024-pixel patches were extracted at 40x magnification for evaluation of both the *NBT-Classifier* and HistoROI. The dataset comprised 8,000 patches per class across the following categories: normal (peri-tumoral, N), benign (pathological benign, PB; usual ductal hyperplasia, UDH), precancerous (flat epithelial atypia, FEA; atypical ductal hyperplasia, ADH), and cancerous (ductal carcinoma in situ, DCIS; invasive carcinoma, IC). Of these, 96.14% were predicted as epithelium by the *NBT-Classifier*, indicating good generalisability on external datasets. Notably, the epithelium probability distributions were shifted for normal breast tissue compared with pathological categories. In contrast, the mixed-tissue-based HistoROI exhibited substantial overlap between the two classes (**Fig. 3b**). ROC analysis further demonstrates that the *NBT-Classifier* outperforms HistoROI in discriminating normal breast epithelium from abnormal tissue (**Fig. 3c**, DeLong’s test, P < 0.0001). Importantly, differences between normal and precancerous or cancerous epithelium were consistently better captured by the *NBT-Classifier* (**Fig. 3d**). When applying Grad-CAM to patches of normal epithelium, we observed differences in the resolution and granularity of attention due to the distinct CNN architectures used (MobileNet^44^ for *NBT-Classifiers* and ResNet18^37^ for HistoROI). Beyond these architectural differences, HistoROI showed limited recognition of entire normal lobules, with attention misaligned and incomplete, whereas the normal-specific *NBT-Classifier* provided more comprehensive and well-aligned coverage of normal epithelium (**Fig. 3e**). Together, these findings suggest that the *NBT-Classifier* captures normal-specific features, in contrast to the more general epithelial features learned by models trained on mixed-breast-tissue datasets.

### Slide-level tissue compartment classification and visualisation

*NBT-Classifiers* are patch-level tissue classification models that, once trained on annotated patches, can be applied to entire WSIs. To demonstrate this, we performed whole-slide tissue classification on a representative WSI using both the 512px- and 1024px-based *NBT-Classifiers*. The results were then converted and imported into QuPath v0.3.0 for interactive histological inspection (**Fig. 4a**, **4b**, **Methods**). Heatmaps generated by both models showed strong alignment between predicted tissue classes and the underlying histology at different scales. For a larger tissue region (5120 × 5120 pixels, 0.25 µm/pixel), we created a smoother heatmap by using predicted tissue class probabilities (**Methods**). This high-resolution heatmap effectively delineated individual lobules and segmented tissue compartments with smooth transitions (**Fig. 4c**). Additionally, we analysed six other regions from tissue across different patient age groups, all of which demonstrated similarly strong performance (**Supplementary Fig. 19**). These results highlight the potential of this approach as a foundation for more advanced downstream image analyses, such as lobule detection.

**Figure 4.**
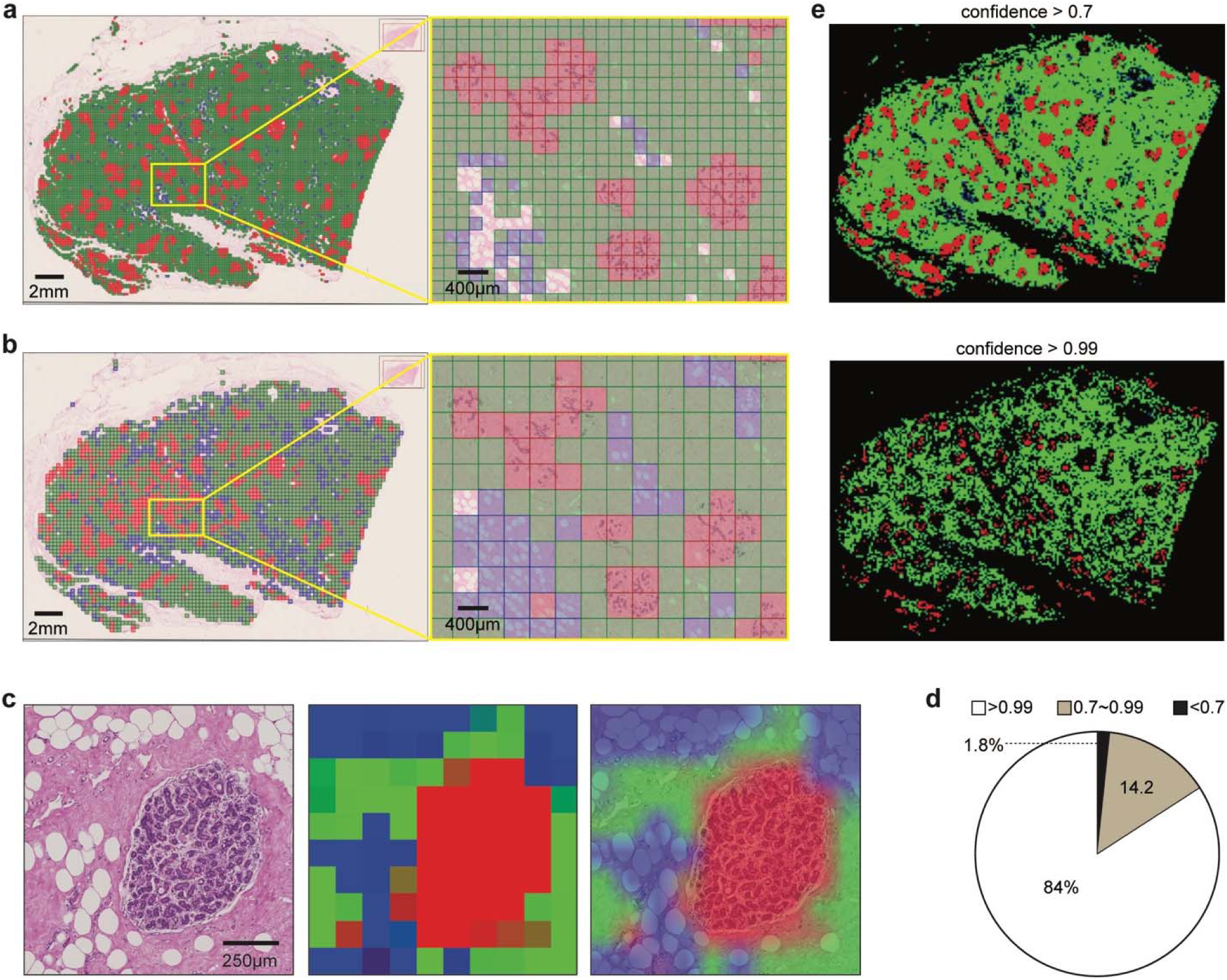
Visualisation of whole slide tissue classification. **a-b** QuPath v0.3.0 visualisation of whole slide tissue class predictions by the 512px- (**a**) and 1024-based (**b**) *NBT-Classifiers*. The original tissue class predictions of the entire WSI were converted into JSON files and imported into QuPath v0.3.0 as pseudo patch-level annotations, shown in the right images. Epithelium, stroma, and adipocytes are represented in red, green, and blue, respectively. Scale bar, 2 mm. Magnified regions are displayed on the right. Scale bar, 400 µm. **c** Tissue probability heatmaps of a representative region. A 5120 × 5120-pixel region containing mixed tissue components (left) was selected and tessellated into non-overlapping 512 × 512-pixel patches. For each patch, the probabilities of epithelium, stroma, and adipocytes were predicted using the 512px-based *NBT-Classifier* and mapped to the red (R), green (G), and blue (B) channels, respectively, to generate a tissue probability heatmap (middle). The heatmap was then resized and overlaid onto the H&E image to facilitate comparison with the corresponding histological patterns (right). Scale bar, 250 µm. **d** Pie chart illustrating the distribution of the 512px-based *NBT-Classifier’s* confidence in the predicted tissue class. **e** Visualisation of patches predicted with varying confidence. The top image shows patches classified with a confidence higher than 0.7, while the bottom image shows patches classified with a confidence higher than 0.99.

Among all patches used for external validation, 98.2% had probabilities greater than 0.7, and 84% had probabilities greater than 0.99 (**Fig. 4d**). We then examined the spatial distribution of patches predicted with varying confidence in the same WSI example. Patches with confidence above 0.7 covered most tissue regions, while those with scores above 0.99 covered only partial regions, with gaps primarily at the boundaries between tissue compartments and within large lobules (**Fig. 4e**). These low-confidence patches in large lobules may explain the reduced accuracy of the 224px-based *NBT-Classifier*, as the smaller patch size tends to capture more intra-lobular stroma, leading to a higher rate of misclassification of epithelium as stroma. Furthermore, the varying spatial localisation of model confidence could serve as a valuable metric for capturing spatial and histological variations within each tissue compartment, making it particularly useful for patch selection in slide-level analysis, such as multiple-instance learning (MIL)^16,45^.

### End-to-end pipeline for analysing large-scale WSIs of NBT

Building on the demonstrated utility of the proposed *NBT-Classifiers*, we integrated the models into an end-to-end WSI pre-processing pipeline (**Fig. 5a**). This generates tissue classification results that facilitate large-scale digital image analysis within target sub-tissue regions in the normal breast, such as lobules and their microenvironment, and supports the patch selection process for more advanced deep learning frameworks, such as MIL^16,45^. The pipeline begins with HistoQC^46^ to detect foreground tissue regions (**Supplementary Fig. 20**, **Methods**). Once identified, the region is tessellated into non-overlapping patches, which are then analysed by the corresponding *NBT-Classifier* to predict tissue classes. This produces a whole slide tissue class heatmap, from which lobules and their peri-lobular regions within a specified range can be detected and localised (**Fig. 5b**, **Methods**). For frameworks such as MIL^16,45^, patches from target tissue regions can be selectively balanced and treated as a hyperparameter to optimise performance during training (**Fig. 5a**). Besides, the pipeline outputs a binary mask that localises individual lobules on each WSI and can be directly imported into QuPath v0.3.0 as pseudo lobule annotations (**Fig. 5c**). Within each lobule, QuPath v0.3.0’s built-in nuclei detection and advanced spatial analysis tools, such as Delaunay triangulation, can be utilised for object-level lobule quantification. Additionally, the original patch-level classifications of the target regions (lobules and peri-lobular areas) can be exported and loaded into QuPath v0.3.0 where digital image analyses, such as texture analysis can be performed, across the slide at the patch level (**Fig. 5d**). These approaches build upon the previous efforts of standardised quantifications of TDLUs^17–21^, aiming at expanding interpretation of lobules as well as their direct microenvironment, which might unlock novel biomarkers that are indicative of breast cancer precursors. The pipeline performed robustly across WSIs with varying staining at both 40x and 20x magnification (**Supplementary Fig. 21**), offering flexibility for studies involving heterogeneous datasets. The source code running this pipeline is available at: https://github.com/cancerbioinformatics/NBT-Classifier and https://hub.docker.com/repository/docker/siyuan726/nbtclassifier.

**Figure 5.**
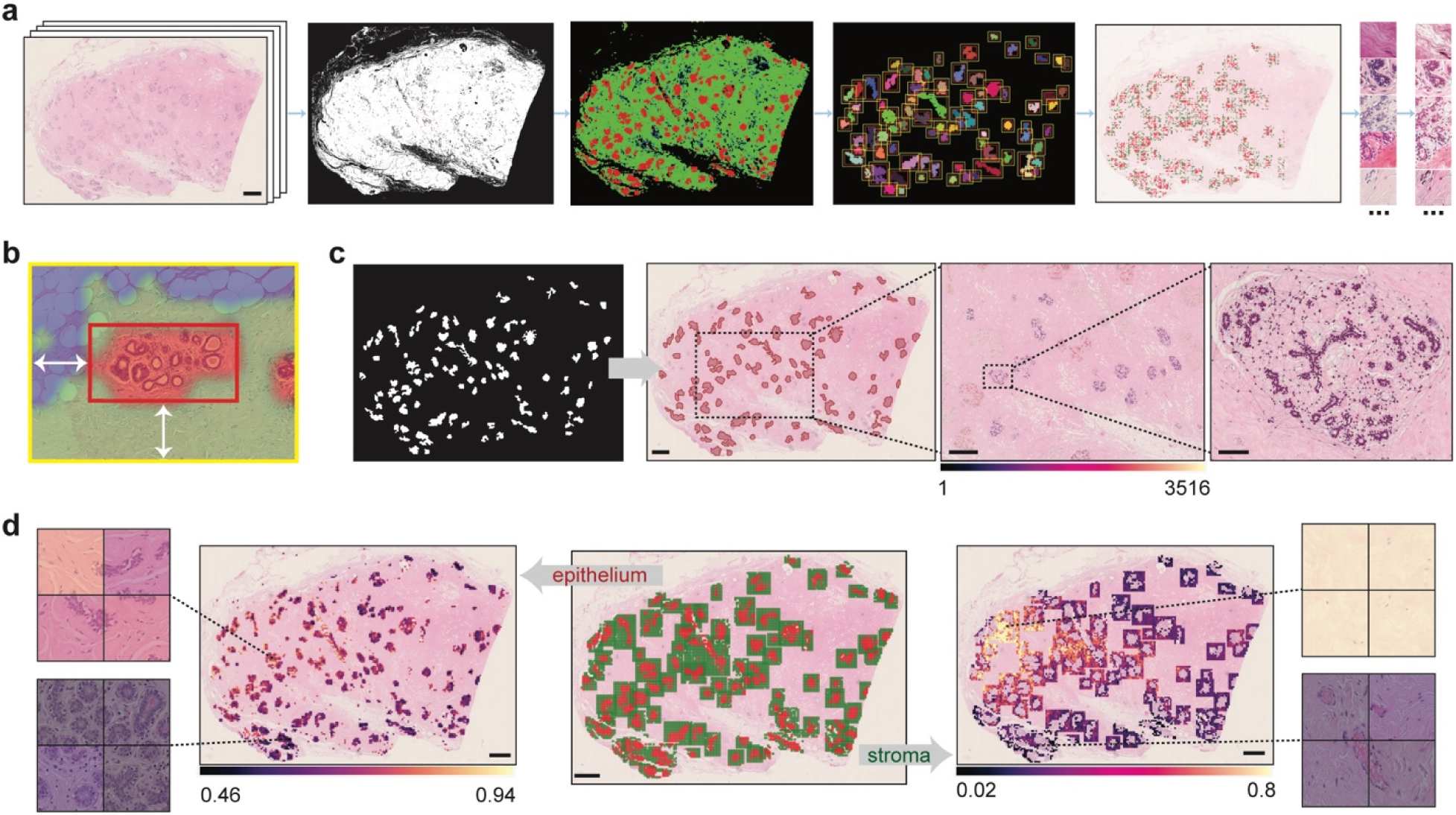
Approaches facilitating downstream WSI analysis of NBTs. **a** Proposed WSI pre-processing pipeline for NBTs. From left to right, the images depict an input WSI (scale bar, 2 mm), detected foreground tissue regions, whole slide tissue classifications (red: epithelium, green: stroma, blue: adipocytes), identified lobular regions, balanced-sampled patches, and stain-normalised patches. **b** Magnified example of a detected lobular region. The tissue class mask was generated by the 512px-based *NBT-Classifier*, resized, and overlaid onto the H&E image. The red box highlights the detected lobule, while the yellow box outlines the lobular region, with a 250 µm (adjustable) width between the two boxes, as indicated by the white arrows. **c** Example demonstrating generated pseudo lobule annotations and downstream nuclei spatial analysis in QuPath v0.3.0. The first image displays a binary mask of detected lobules, which were imported as pseudo-lobule annotations (highlighted in red) in QuPath v0.3.0, as shown in the second image. Scale bar, 2 mm. Within QuPath v0.3.0, nuclei segmentation was performed using StarDist, followed by Delaunay spatial analysis to quantify various graph properties of each lobule, such as the number of nuclei within the graph. The third image visualises the variation in the number of nuclei (indicated by the colour bar) across Delaunay graphs constructed for lobules within a magnified region. Scale bar, 1 mm. The last image presents an example of a Delaunay graph constructed from a magnified lobule. Scale bar, 100 µm. **d** Example of pseudo patch-level tissue class annotations within detected lobular regions and downstream texture analysis in QuPath v0.3.0. The middle image displays pseudo patch-level annotations of the epithelium (red) and peri-lobular stroma (green) in QuPath v0.3.0, which were subsequently analysed separately using Haralick texture features. The left image visualises the variation in the Inverse Difference Moment (F4) across epithelium patches, while the right image shows the variation in the Angular Second Moment (F0) across stroma patches. Scale bar, 2 mm. Magnified patch examples with low and high F4 and F0 values are displayed on the side.

## Discussion

AI-driven WSI analysis has given rise to groundbreaking advancements in histopathology^13^. In breast research, while the research frontier remains largely focused on breast cancer, histologically normal tissues are significantly underrepresented among millions of digitised WSIs, with few applications specifically designed for facilitating in-depth morphological characterisation of NBTs^17–24^. To address this gap, we compiled high-quality, manually annotated WSIs of NBTs, sourced from women with varying breast cancer risk across multiple institutions. Through extensive optimisation and external validation, we developed robust *NBT-Classifiers* capable of processing WSIs at different scales and performing patch-level tissue classification of epithelium, stroma, and adipocytes. To the best of our knowledge, *NBT-Classifiers* represent the first deep learning-based CPath models specifically trained to classify tissue compartments within normal breast tissue (from individuals without any lesions). Building on this, we integrated an end-to-end WSI pre-processing pipeline to ensure objective and reproducible tessellation and patch selection from biologically important regions, such as lobules and their microenvironment. The outputs are compatible with QuPath v0.3.0, one of the most widely used digital image analysis platforms and can be seamlessly integrated with deep learning-based CPath frameworks such as MIL^16,45^. Our models (https://github.com/cancerbioinformatics/NBT-Classifier and https://hub.docker.com/repository/docker/siyuan726/nbtclassifier) and datasets (https://github.com/cancerbioinformatics/OASIS) are publicly available, whereby the latter will be a highly valuable resource for training pathology foundation models as they provide rich histology data of normal breast.

To establish fundamental tissue classification tools for NBTs, particularly as a primary step for downstream analyses, ensuring generalisability, robustness, and adaptability is essential. Achieving this requires a diverse and representative WSI dataset, ground-truth manual annotations, and seamless compatibility with downstream analytical frameworks. Previous NBT studies have typically relied on a single data source, such as benign breast biopsies^17–19,23^, healthy donors^20,22^, or reduction mammoplasties^21^. Only one exception^24^ incorporated NBTs from both risk-reducing mastectomy and reduction mammoplasty specimens (*gBRCA1/2m* status unknown). However, these datasets do not capture the diverse origins of NBTs, which may exhibit varying predispositions to cancer initiation and distinct histological manifestations. In contrast, a key strength of our work is the incorporation of a broad spectrum of NBT variations, including differences in patient age, breast cancer risk, H&E staining protocols, and scanning platforms.

In the generation of AI, manual annotations should not be overlooked when developing supervised applications, especially in highly expert histopathological domains^47^. Our fully pathologist-supervised ground-truth likely explains the slightly better performance in stroma and adipocytes classification compared to HistoROI^16^, which employed human-in-the-loop annotations. Among the previous NBT research, three studies provide manual annotations, yet their sample sizes and annotation levels vary (**Supplementary Table 4**). We expanded on this by providing detailed annotation of lobules, stroma and adipocytes that displayed biological variations. Moreover, we demonstrated that *NBT-Classifiers*, trained exclusively on histologically normal breast tissue, learn distinctive, normal-specific features. Their outputs reflect the degree of similarity to truly normal epithelium, whereas models trained on mixed breast tissues capture more general epithelium features. The comparable performance observed in the mix-breast-tissue model might largely arise from the relative similarity of epithelium to other tissue compartments, making the prediction effectively an approximation rather than a normal-specific assessment. This distinction is critical, as a classifier that is grounded in the unique characteristics of normal breast tissue offers a more robust foundation for future applications in early detection, positioning it as a screening tool for identifying pathological deviations in gigapixel WSIs of histologically normal breast tissue.

Performing patch-level analysis within key regions of NBTs, such as lobules and their microenvironment, is essential for detecting early cancer precursors. Prior NBT studies have focused on well-established cancer risk biomarkers, such as TDLU involution^17–20^, immune infiltration in lobular regions^21^, alterations in tissue composition^22,23^, and fibroglandular density^24^. In addition to these visible pathological biomarkers, recent studies have highlighted greater predictive potential within NBTs at the single-cell level^48–51^. Given the extensive validation of genotype-phenotype links in various CPath studies focused on cancer^52–54^, analysing sub-tissue regions at the cellular and sub-cellular level shows significant promise for NBTs as well. In this context, the ability to adapt to broader downstream pathomics feature extraction tools, such as QuPath-based digital image analysis, or deep learning-driven CPath frameworks, opens new opportunities for evaluating high-throughput histopathological measurements. This, in turn, provides insights into localised histological markers that may signal the earliest stages of breast cancer initiation. Building upon this, advanced techniques such as spatial transcriptomics^55^ and spatial proteomics^56^ could uncover the underlying molecular mechanisms driving these histological changes. By linking molecular data with histological features, a deeper understanding of how early molecular alterations contribute to the development of breast cancer can be achieved.

One limitation of this study is the diversity of the external validation cohorts, which may not fully capture the spectrum of variations in NBTs, such as contralateral tissues or those from risk-reducing mastectomies. Future work will systematically evaluate the model’s performance particularly across diverse demographic groups and healthcare settings, with a specific focus on ensuring equitable generalisability in underrepresented populations. Another limitation lies in the annotation scope, which included only subsets of tissue types. Consequently, regions like artefacts, necrosis and blood vessels were misclassified as epithelium when applying the *NBT-Classifiers* to whole WSIs. Including these regions as additional classes could improve classification accuracy. While expert annotation remains essential due to the histological complexity of NBTs, strategies to improve efficiency, such as AI-powered annotation platforms or carefully guided crowdsourcing approaches^47^, could help scale the process. Biological assays or molecular markers^21^ could further support the generation of more precise, cell-level ground-truth annotations. In addition, ViT-based foundation models were not evaluated, as the near-perfect AUCs achieved suggest that the semantic distinctions between epithelium, stroma, and adipocytes were sufficiently pronounced for capture by conventional CNNs. Additionally, while ViTs excel at capturing global context, their patch-based architecture and multi-head attention often produce coarse or fragmented saliency maps, limiting interpretability at the cellular and sub-tissue level^57,58^. In contrast, CNNs, with their locality and translation-invariance biases, generate more granular and biologically intuitive explanations that align with pathologists’ reasoning^59^. Thus, although CNNs may lack some global context, their explainability was critical for our goal of visualising distinct, plausible patterns in normal breast tissue. Future work could explore whether transformer-based backbones provide additional performance gains or improve generalisability. Lastly, due to out-of-distribution effects between training and testing data, domain shift was observed in features extracted from both *NBT-Classifiers*, as well as HistoROI (**Supplementary Fig. 22**), suggesting stain normalisation cannot fully address the variations in data distributions across cohorts. Recent studies indicate that even foundation models^60,61^ trained on large and diverse datasets do not always avoid domain shift^62,63^. In future work, domain generalisation techniques could be applied to improve the models, enabling them to learn domain-invariant features and reduce cohort-specific biases^30,64^.

In summary, we present deep learning-based *NBT-Classifiers* for patch-level tissue type classification in NBTs. These have the potential to enhance our understanding of how various NBT components contribute to both benign and malignant breast pathology and lay the groundwork for the development of more advanced deep learning models and spatial-defined molecular large-scale analyses in the future.

## Methods

### Patient cohorts

In this study, NBTs were obtained from the following sources: core biopsies of healthy donors (non-*gBRCA1/2m* carriers, n=12 WSIs); women undergoing reduction mammoplasties (non-*gBRCA1/2m* carriers, n=30 WSIs); *gBRCA1/2m* carriers undergoing risk-reducing mastectomy (n=21 WSIs); and contralateral or ipsilateral normal tissues from breast cancer patients (n=7 WSIs). A total of 70 digitised formalin-fixed, paraffin-embedded (FFPE), haematoxylin and eosin (H&E)-stained WSI were collected across five cohorts: the King’s Health Partners Cancer Biobank (KHP) in London (UK) (n=16 WSIs), the Netherlands Cancer Institute (NKI) in Amsterdam (Netherlands) (n=16 WSIs), the Barts Cancer Institute (BCI) in London (UK) (n=16 WSIs), the École Polytechnique Fédérale de Lausanne (EPFL) in Lausanne (Switzerland) (n=10 WSIs), and the public Susan G. Komen Tissue Bank (SGK) (n=12 WSIs)^29^. Patient age ranged from 16 to 74 years. WSIs from the KHP, BCI, and EPFL cohorts were scanned on Hamamatsu NanoZoomer scanners whilst WSIs from the NKI cohort were scanned on MIRAX scanners, all at 40x magnification (0.25 µm/pixel). WSIs from the SGK cohort were downloaded from the virtual tissue bank (https://virtualtissuebank.iu.edu/query/), having been scanned at 20x magnification^29^.

To develop *NBT-Classifiers*, we used the NKI and BCI cohorts. These included samples of NBTs from reduction mammoplasties (n=4 patients, non-*gBRCA1/2m* carriers), risk-reducing mastectomies in *gBRCA1/2m* mutation carriers (n=21 patients, no cancer at surgery), and contralateral or ipsilateral NBTs from breast cancer patients (n=7 patients). The remaining WSIs from the KHP, EPFL and SGK cohorts were used for external validation. These included NBTs from reduction mammoplasties (n=26 patients, non-*gBRCA1/2m* carriers) and core biopsies from healthy donors (n=12 patients, non-*gBRCA1/2m* carriers). Detailed information on individual slides can be found in **Supplementary Table 1** and is available at the OASIS repository (https://github.com/cancerbioinformatics/OASIS).

### Manual annotation

Expert manual annotations of epithelium, stroma, and adipocytes were performed for each WSI under the supervision of a consultant pathologist (SEP) using QuPath v0.3.0^10^. To improve the efficiency of the annotation process, we adopted a hybrid strategy: lobular boundaries were precisely delineated to annotate epithelium, while stroma and adipocytes were annotated using rectangular boxes of comparable size, randomly sampled from various regions throughout the NBT, to ensure a balanced representation of tissue classes during training (**Fig. 1c, 1d**, **Supplementary Fig. 1**). The pathology-guided annotated WSIs of NBTs are available in the nOrmal breASt tISsue Dataset (OASIS) repository, accessible at https://github.com/cancerbioinformatics/OASIS.

### Patch-level datasets for training

To train models classifying NBTs at different scales, we used QuPath v0.3.0 to extract labelled patches from the NKI and BCI cohorts. These datasets consisted of non-overlapping patches with sizes of 224 × 224, 512 × 512, and 1024 × 1024 pixels, all extracted at a resolution of 0.25 µm/pixel (**Supplementary Table 2**). Only patches with their centres located within the annotated regions were extracted and inherited the labels from the pixel-level manual annotations. The patch extraction was performed using Groovy scripts in QuPath v0.3.0. The script is available at: https://github.com/cancerbioinformatics/NBT-Classifier/blob/main/patching_qupath.groovy.

### Stain normalisation

In experiments optimising the stain normalisation method, we used the same reference image when implementing the Reinhard, Macenko, and Vahadane method, which is available at: https://github.com/cancerbioinformatics/NBT-Classifier/blob/main/data/he.jpg.

### Architecture of NBT-Classifiers

The *NBT-Classifiers* leverage transfer learning, each employing a pre-trained CNN backbone to extract visual features from image patches, followed by a trainable classification head for tissue class prediction (**Supplementary Fig. 4**). Specifically, the feature maps from the last convolutional layer of each CNN backbone were transformed into a single-dimensional vector using a global average pooling layer. The reduced visual features serve as input to the subsequent classification module, which consists of two densely connected layers. The first dense layer linearly transforms the input features into a 1024-dimensional representation, followed by a second dense layer that reduces it to 512 dimensions, both employing the “ReLU” activation function. To mitigate overfitting, a Dropout layer with a rate of 50% is applied between these layers. The final output layer maps the 512-dimensional representation to a 3-dimensional feature, using a “Softmax” activation function to predict the probability distribution across the three tissue classes. The class with the highest probability is assigned as the predicted tissue class (0: epithelium, 1: stroma, 2: adipocytes).

### Three-fold cross-validation

*NBT-Classifiers* were trained and optimised through three-fold cross-validation. To ensure methodological consistency and fair comparison across experiments, a single fixed partition of the data was applied throughout all analyses. In each fold, patches were split at the WSI level: 20 WSIs for training (10 from each cohort), 4 WSIs for internal validation (2 from each cohort), and 8 WSIs for internal testing (4 from each cohort).

For experiments optimising the stain normalisation method and CNN backbone, patch-level datasets containing non-overlapping 512 × 512-pixel patches from the NKI and BCI cohorts were used. To ensure class balance, 250 patches per tissue class were randomly sampled from each WSI. For experiments optimising *NBT-Classifiers* on patches of different FOVs, non-overlapping patches of 224 × 224, 512 × 512, and 1024 × 1024 pixels from the NKI and BCI cohorts were used. To ensure class balance, 150 patches per tissue class were randomly sampled from each WSI (**Supplementary Fig. 6-7**, **Supplementary Table 2**).

For training patches, six data augmentation techniques were employed, including horizontal flipping, random rotation within the range of -40 degrees to 40 degrees, random shifts, random shear transformations, random zoom in/out, and random shifts of the channel values within the range of [0, 10], to enhance model’s robustness and regularise the algorithm. The weights of the CNN backbones were all frozen, and the densely connected layers in the classification head were initialised by the Xavier normal initializer and set to be trainable. We used categorical cross-entropy as the loss function and Adam optimizer with a learning rate of 1e-5. The batch size was 16 and we trained each model for 50 epochs. The best model was selected according to the highest validation accuracy, reflecting the overall multi-class classification performance, and was then saved for further testing.

### External validation

For cohorts used for external validation (KHP, EPFL and SGK), we extracted non-overlapping patches with a size of 512 × 512 and 1024 × 1024 pixels at a resolution of 0.25 µm/pixel for the KHP and EPFL cohorts. For the SGK cohort, scanned at 20x magnification, we extracted non-overlapping patches with a size of 256 × 256 and 512 × 512 pixels at a resolution of 0.5 µm/pixel (**Supplementary Table 2)**. All patches were normalised using the Reinhard method and patches from SGK were resized to match the dimension of the corresponding *NBT-Classifier*. For each patch, each *NBT-Classifier* outputs probabilities of epithelium, stroma and adipocytes and the tissue class with the highest predicted probability was assigned as the final class prediction. The primary statistical endpoint was the area under the receiver operating characteristic curve (AUROC). The secondary statistical endpoint was class-specific accuracy.

Accuracy was determined by comparing the ground-truth labels with the class assigned the highest predicted probability. We report both the overall accuracy as well as individual accuracies in each KHP, EPFL and SGK cohort.

### Confounder analysis

We evaluated whether the *NBT-Classifiers* exhibited any bias towards potential confounders such as patient age and the source of NBTs. For a fair comparison, we sampled 15 patients from a total of 38 patients across three external cohorts, ensuring balanced representation across three age groups, namely “premenopausal years” <45 years: 5 patients; “menopausal years” 45–55 years: 5 patients; “postmenopausal years” >55 years: 5 patients. We sampled 30 patients from a total of 32 patients across NKI and BCI cohorts to include equal representation across three NBT sources: reduction mammoplasty (10 patients), risk-reducing mastectomy (10 patients), and NBTs from breast cancer patients (10 patients). To mitigate class imbalance, 500 patches per tissue class per WSI were randomly sampled when using the 512px-based model and 250 patches per class per WSI were sampled when using the 1024px-based model. These experiments were repeated three times.

### Implementation of HistoROI

HistoROI^16^ is a CNN-based tissue classification model trained on WSIs of histologically normal, noncancerous, precancerous, and cancerous breast tissue from the public BReAst Carcinoma Subtyping (BRACS) dataset (https://research.ibm.com/haifa/Workshops/BRIGHT/, https://www.bracs.icar.cnr.it/)^42,43^. The model classifies epithelium, stroma, and adipocytes, along with lymphocytes, miscellaneous tissues, and artefacts. It processes 256 × 256-pixel patches at a resolution of 1 µm/pixel, capturing the same field of view (FOV) as 1024 × 1024-pixel patches at 0.25 µm/pixel. To ensure a fair comparison, we resized the 1024 × 1024-pixel patches extracted from WSIs in three external cohorts and applied Reinhard stain normalisation before using them with HistoROI^16^.

### Analysis of epithelium patches from BReAst Carcinoma Subtyping (BRACS) database and analysis

To evaluate the ability of *NBT-Classifier* and HistoROI to discriminate between histologically normal, precancerous, and cancerous epithelium, we systematically curated a patch dataset from the public BRACS database^42,43^ that was used to develop the HistoROI model. Specifically, region of interest (ROI) images (scanned at 40x, available at: https://www.bracs.icar.cnr.it/) encompassing peri-tumoral normal breast tissue (N), atypical ductal hyperplasia (ADH), ductal carcinoma in situ (DCIS), and invasive carcinoma (IC) were tessellated into patches compatible with both models. We then randomly sampled 8,000 patches per class and applied both classifiers to obtain epithelium probabilities and feature representations. The distributional differences in epithelium probabilities between the normal class (N) and each pathological class (ADH, DCIS, IC) were assessed using the Wilcoxon rank-sum test, followed by Bonferroni correction for multiple comparisons. The code is available at: https://github.com/cancerbioinformatics/NBT-Classifier/blob/main/notebooks/BRACS_analyses.ipynb.

### Visual assessment of feature representations

We performed t-distributed Stochastic Neighbour Embedding (t-SNE)^41^ to visualise the output of features from the layer before the final output layer, of each *NBT-Classifier* in a 2-dimensional space. To facilitate the inspection of the corresponding histology of the features, we randomly selected 500 H&E patches from each ground-truth tissue class (epithelium, stroma, adipocytes) and projected them onto the t-SNE plot based on their coordinates. The code for this visualisation is available at: https://github.com/cancerbioinformatics/NBT-Classifier/blob/main/vis_features.ipynb.

### CAM and Grad-CAM visualisation

To enhance model interpretability, we applied Class activation mapping (CAM)^27^ and Gradient-weighted Class activation mapping (Grad-CAM)^28^. These techniques generate activation heatmaps that highlight the spatial regions most important to the model’s predictions for any given input image, utilising the final convolutional layer of the MobileNet backbone weighted by importance scores. Based on their distinct mechanisms (**Supplementary Fig. 13**), CAM typically generates more localised heatmaps that highlight areas relevant to the final classification. In contrast, Grad-CAM may produce more diffuse or generalised heatmaps in some instances, due to its reliance on gradients to capture the model’s learning dynamics. Once the activation heatmaps were computed, we applied bicubic interpolation to match their dimensions to those of the original H&E images. The resulting smoothed heatmaps were overlaid onto the digitised H&E images to visualise the high-attention histological structures corresponding to each particular tissue class prediction. The source code is available at: https://github.com/cancerbioinformatics/NBT-Classifier/blob/main/vis_CAMs.ipynb.

### Foreground tissue detection

Given that WSIs often contain substantial non-tissue regions, including white background and artefacts such as coverslip edges, excluding these regions is essential for reducing the model’s inference time. We employed HistoQC^46^, a WSI quality control software, to automatically generate binary masks that highlight the artefact-free foreground tissue regions on each WSI (**Supplementary Fig. 20**), ensuring that only tissue-containing regions are included in downstream tissue classification analyses.

### Whole slide tissue classification

When using the 512px-based *NBT-Classifier*, the detected foreground tissue regions are divided into non-overlapping patches with a fixed size of 128 × 128 µm to account for variations in the micron per pixel (mpp) values across different scanners. These patches are then resized to 512 × 512 pixels to match the input dimensions of the model. When using the 1024px-based *NBT-Classifier*, the detected foreground region is divided into non-overlapping patches with a fixed size of 256 × 256 µm and resized to 1024 × 1024 pixels. Each *NBT-Classifier* outputs both probabilities of epithelium, stroma and adipocytes and the final class prediction for each patch, along with the original coordination on the slide.

### Whole slide visualisation of tissue classification results

To visualise the whole slide tissue classification in a single image, we arranged the class predictions of patches according to their corresponding coordinates on the slide to create a heatmap. For interactive histological inspection, the tissue class heatmap can be exported as a JSON file and imported into the QuPath v0.3.0 platform. Additionally, we generated smoother tissue class heatmaps by upscaling the predicted probability matrix through interpolation. The visualisation method is available at: https://github.com/cancerbioinformatics/NBT-Classifier/blob/main/NBT_pipeline.ipynb.

### Lobule detection

To detect individual lobules with their surrounding peri-lobular regions (adjustable), the epithelial layer in the whole slide tissue class heatmap is extracted and enlarged by a factor of 32. Then, connected component analysis is applied to identify groups of connected foreground pixels (filled with a value of 1) as distinct objects. Through manual examination, we observed that lobules empirically are covered by more than two epithelium patches, approximately 500,000 pixels and 0.25 µm/pixel. Therefore, we removed foreground objects (filled with a value of 1) smaller than 400,000 pixels and filled “holes” (filled with a value of 0) smaller than 400,000 pixels inside each detected foreground object. For slides scanned at 20x magnification, the threshold is adjusted to 250,000 pixels. This step reduces noise and increases the robustness for lobule localisation. This method outputs a binary mask highlighting individual lobules and an optional surrounding stroma within a pre-defined range. The mask can be converted into a JSON file and imported into QuPath v0.3.0 for downstream digital image analyses. The source code is available at: https://github.com/cancerbioinformatics/NBT-Classifier/blob/main/NBT_pipeline.ipynb.

### Deep learning implementation

All deep learning experiments were implemented with TensorFlow2 in Python on two NVIDIA A100 GPUs from the high-performance computing cluster of King’s Computational Research, Engineering, and Technology Environment (CREATE).

### Statistical analysis

To compare distributions between two groups, we performed pairwise Wilcoxon tests, applying Bonferroni correction for multiple comparisons. A p-value of < 0.05 was considered statistically significant. DeLong’s test was employed to compare the AUCs between different classifiers, assessing the significance of performance differences^65,66^.

## Data availability

WSIs involved in this study are stored at the OASIS repository: https://github.com/cancerbioinformatics/OASIS, which currently can be accessed upon request.

## Code availability

The implementation code involved in this study can be found at: https://github.com/cancerbioinformatics/NBT-Classifier, including Python codes for implementing *NBT-Classifiers* and the pre-processing pipeline, and QuPath v0.3.0 scripts for tessellation and importing pseudo annotations.

## Acknowledgements

The authors wish to acknowledge the support of the King’s Health Partners Cancer Biobank in London, the Netherlands Cancer Institute in Amsterdam, the Barts Cancer Institute in London, and the École Polytechnique Fédérale de Lausanne in Lausanne, Switzerland and Aasiyah Oozeer, Marcello D’Angelo, Rachel Barrow, Rachel Nelan, Marcelo Sobral-Leite, Fabio de Martino for material collection.

The authors would like to thank all members of the Cancer Bioinformatics team at King’s College London (London, UK) for their helpful suggestions. We thank the Breast Cancer Research Trust, Breast Cancer Now (and their legacy charity Breakthrough Breast Cancer), the Medical Research Council (MRC) [MR/X012476/1], Cancer Research UK [CRUK/07/012, KCL-BCN-Q3], CRUK City of London Centre Award [CTRQQR-2021/100004], and the UK Government through the Research Venture Catalyst award, Department for Science, Innovation, and Technology for funding this project. Siyuan Chen is funded by a China Scholarship Council PhD scholarship. The funders had no role in study design, data collection and analysis, decision to publish, or preparation of the manuscript. During the preparation of this work, the author(s) used Grammarly (free version) and ChatGPT (version 2) to correct some grammatical errors and enhance the overall readability of the manuscript. After using this tool/service, the authors reviewed and edited the content as needed and take(s) full responsibility for the content of the publication.

## Author Contributions

**Conceptualisation**: Anita Grigoriadis, Sarah Pinder, Siyuan Chen, Mario Parreno-Centeno, Christopher R.S. Banerji. **Data curation**: Siyuan Chen, Graham Booker. **Formal analysis**: Siyuan Chen, Mario Parreno-Centeno. **Funding acquisition**: Anita Grigoriadis. **Investigation**: Anita Grigoriadis, Sarah Pinder, Siyuan Chen, Mario Parreno-Centeno, Christopher R.S. Banerji. **Methodology**: Siyuan Chen, Mario Parreno-Centeno, Christopher R.S. Banerji, Gregory Verghese, Salim Arslan. **Project administration**: Anita Grigoriadis. **Resources**: Cheryl Gillett, Louise J. Jones, Esther H. Lips, Cathrin Brisken, Aasiyah Oozeer, Marcello D’Angelo, Rachel Barrow, Rachel Nelan, Marcelo Sobral-Leite, Fabio de Martino. **Software**: Siyuan Chen, Mario Parreno-Centeno. **Supervision**: Anita Grigoriadis, Sarah Pinder, Christopher R.S. Banerji. **Validation**: Fathima Sumayya Mohamed. **Visualisation**: Siyuan Chen. **Writing – original draft**: Siyuan Chen. **Writing– review & editing**: Anita Grigoriadis, Sarah Pinder, Christopher R.S. Banerji, Esther H. Lips, Cathrin Brisken, Louise J. Jones, Matthew J. Smalley, Gregory Verghese, Fathima Sumayya Mohamed, Graham Booker, Siyuan Chen, Mario Parreno-Centeno, Salim Arslan, Pahini Pandya.

## Competing interests

Anita Grigoriadis, Louise J. Jones and Greg Verghese are Co-Founders of PharosAI. Salim Arslan and Pahini Pandya and employed by Panakeia Technology, UK. All other authors declare no relevant conflict of interest.

## Notes

https://github.com/cancerbioinformatics/OASIS

https://github.com/cancerbioinformatics/NBT-Classifier

### Summary of Updates

Section on Training exclusively on normal tissue enables learning of distinctive features in the normal breast updated to clarify NBT-Classifiers capture features unique to normal breast epithelium; Figure 3 revised; Supplemental files updated

