## Supplementary Fig. 1 for "Normal Breast Tissue (NBT)-Classifiers: Advancing Compartment Classification in Normal Breast Histology"

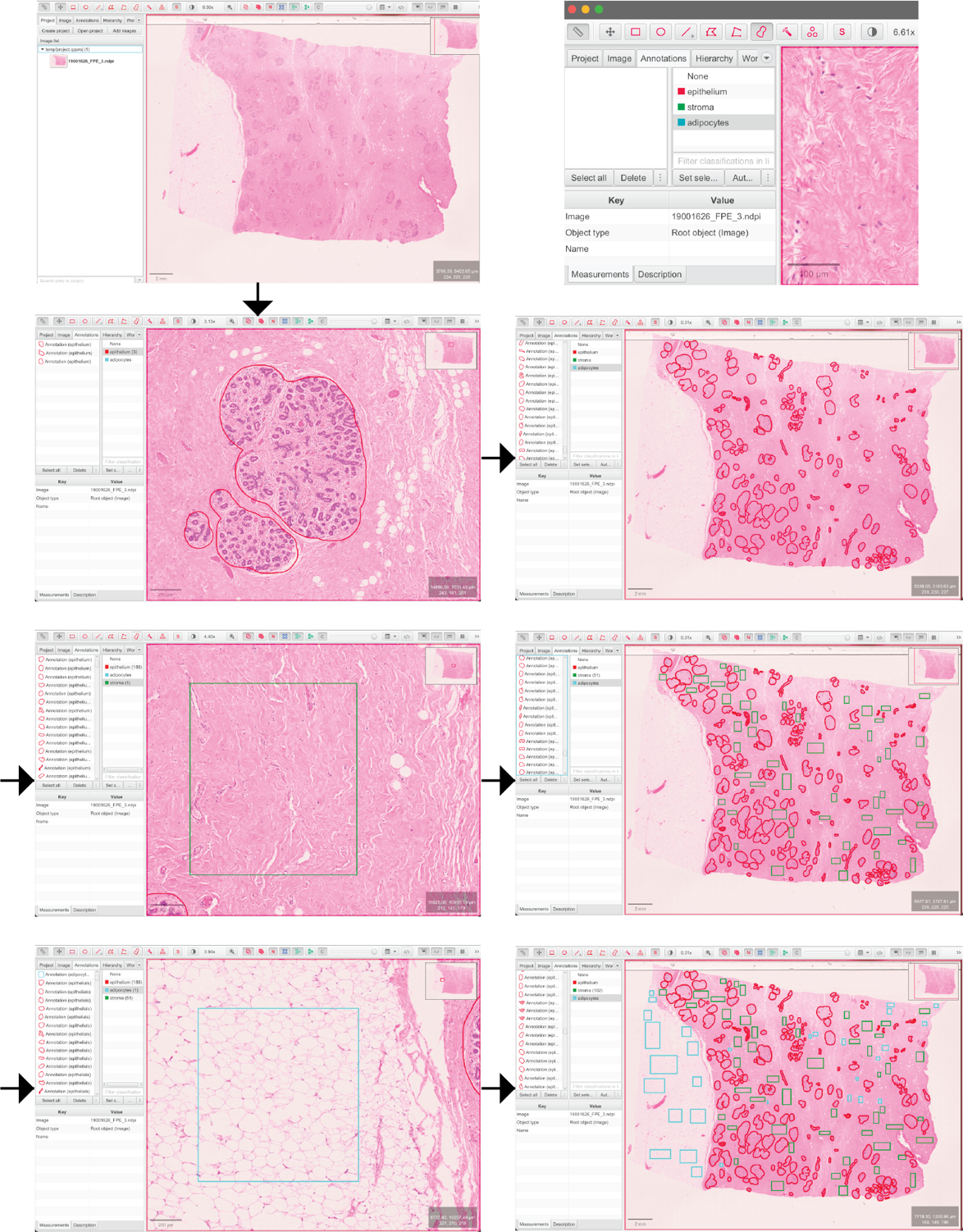

**Supplementary Fig. 1 Schematic pipeline of the manual annotation procedure using QuPath v0.3.0.** For each WSI, epithelium regions (in red) were exhaustively delineated using the brush tool to ensure precise annotation. To obtain a balanced number of training patches from each tissue class, similarly sized regions of stroma (in green) and adipocytes (in blue) were randomly selected from various areas of the WSI using rectangular boxes. This approach enhances efficiency while ensuring the inclusion of spatial variations and maintaining proportional representation across tissue types.
