## Supplementary Fig. 2 for "Normal Breast Tissue (NBT)-Classifiers: Advancing Compartment Classification in Normal Breast Histology"

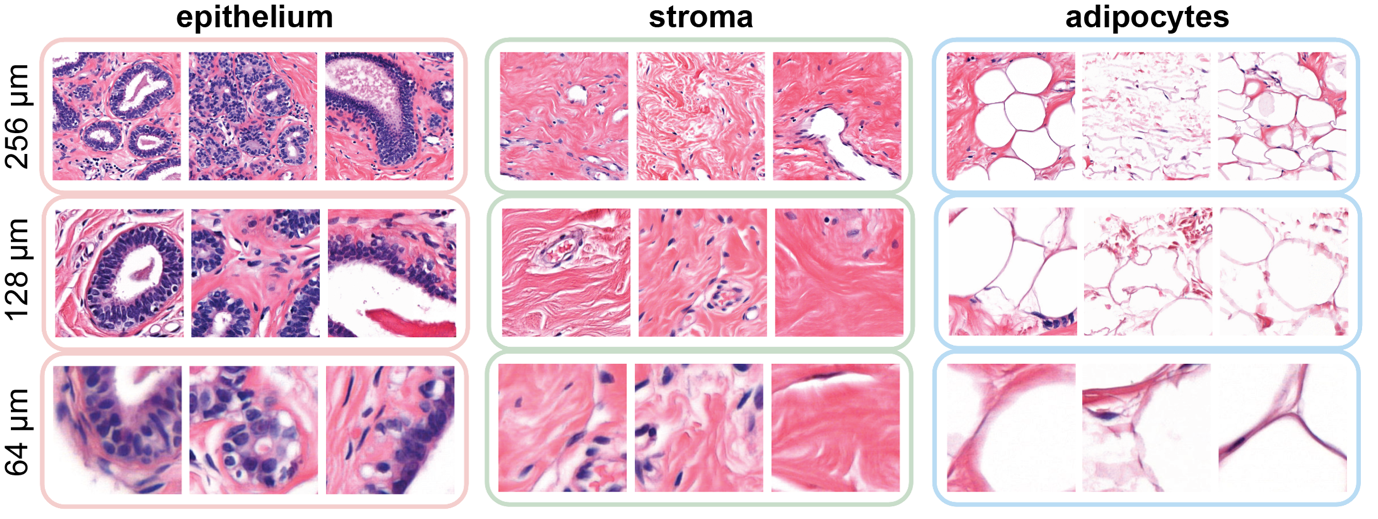

**Supplementary Fig. 2 Illustration of the histology of NBTs captured at different fields of view.** From left to right, the columns display example patches of epithelium, stroma, and adipocytes. Within each column, the rows, from top to bottom, show histological structures captured at fields of view of 256 x 256 µm, 128 x 128 µm, and 64 x 64 µm, respectively.
