## Supplementary Fig. 3 for "Normal Breast Tissue (NBT)-Classifiers: Advancing Compartment Classification in Normal Breast Histology"

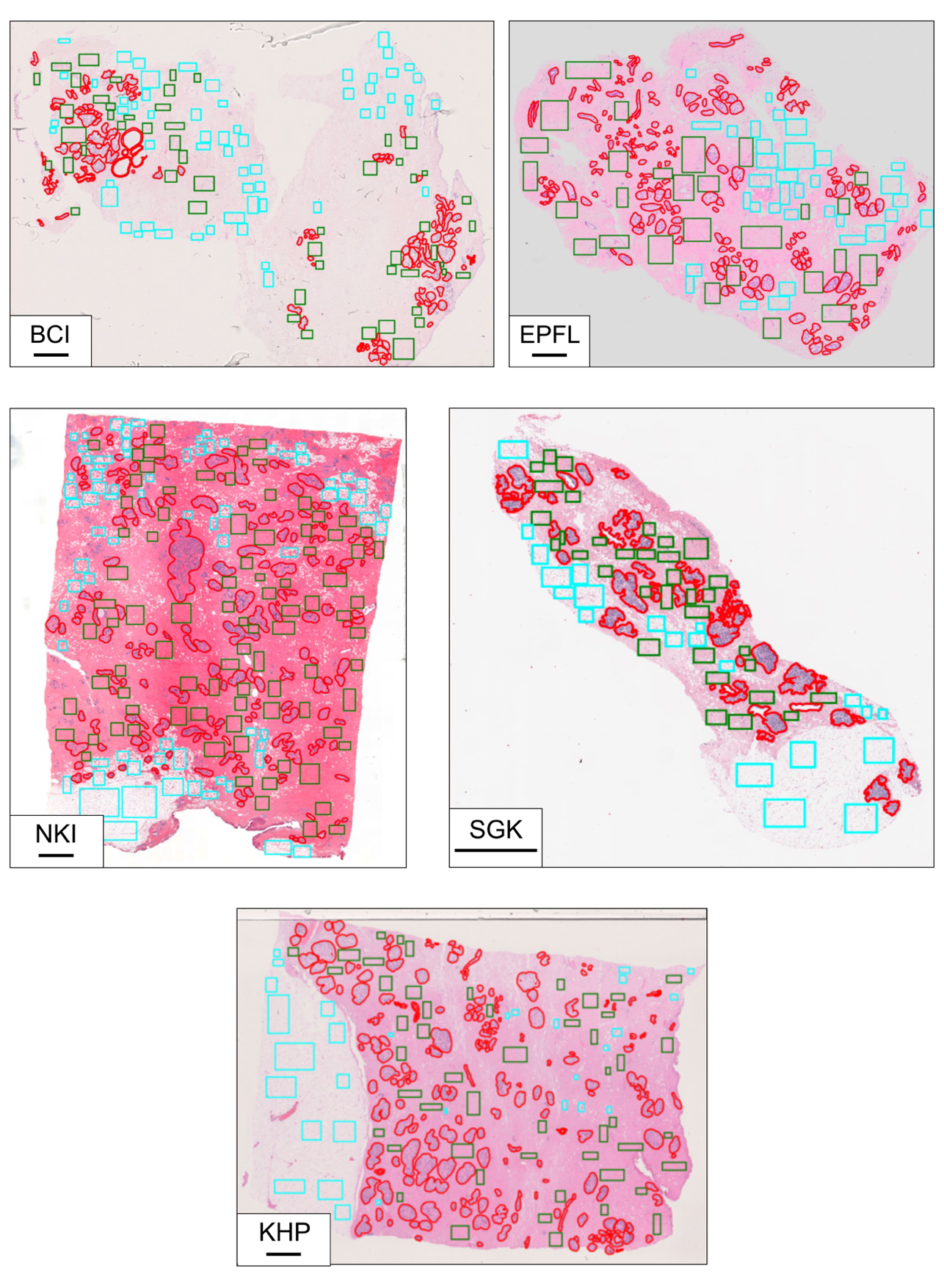

**Supplementary Fig. 3 Examples of annotated WSIs of NBTs from BCI, EPFL, NKI, SGK and KHP cohorts.** Epithelium, stroma, and adipocytes are annotated in red, green, and blue, respectively. Scale bar, 2 mm.
