## Supplementary Fig. 4 for "Normal Breast Tissue (NBT)-Classifiers: Advancing Compartment Classification in Normal Breast Histology"

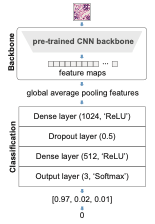

**Supplementary Fig. 4 Illustration of NBT-Classifiers’ architecture.** Each NBT-Classifier utilises a pre-trained CNN backbone to extract feature maps, which are then processed via global average pooling. A classification head subsequently predicts the probabilities for epithelium, stroma, and adipocytes, with the final tissue class assigned based on the highest probability: 0 for epithelium, 1 for stroma, and 2 for adipocytes.
