## Supplementary Fig. 5 for "Normal Breast Tissue (NBT)-Classifiers: Advancing Compartment Classification in Normal Breast Histology"

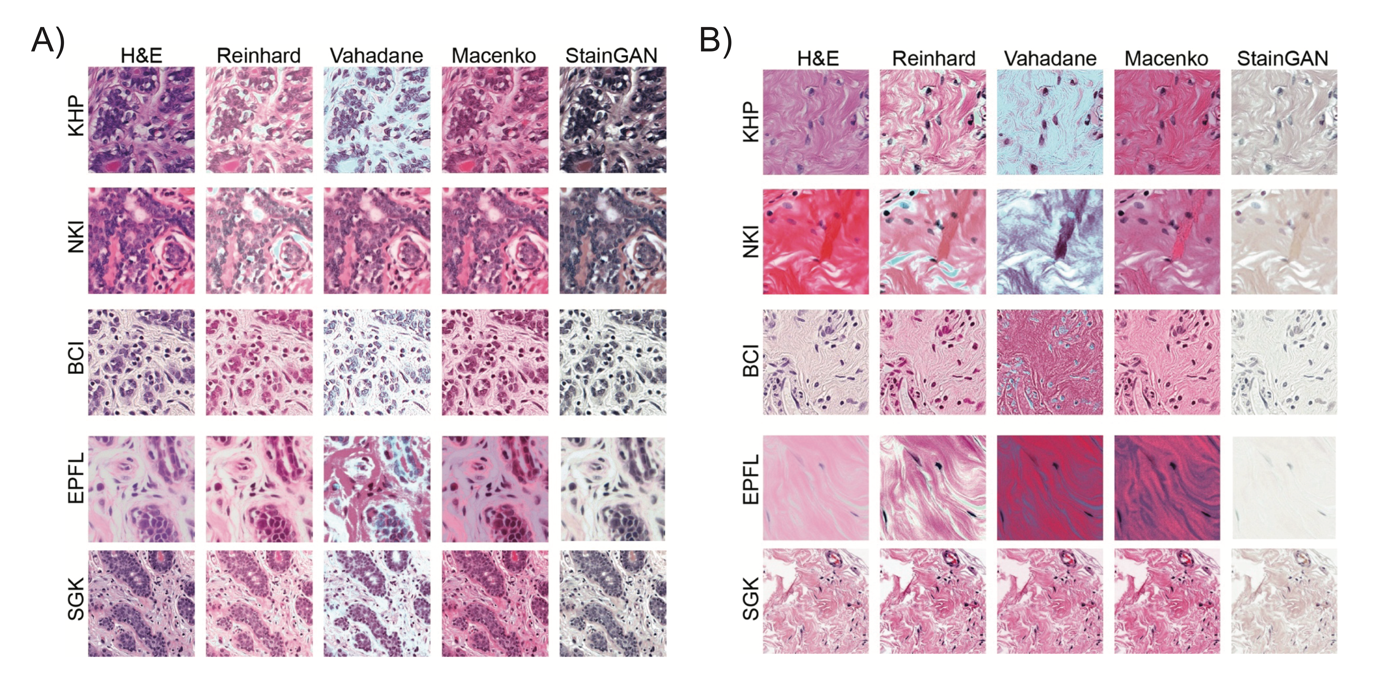

**Supplementary Fig. 5 Illustration of different stain normalisation methods.** The first column shows the original H&E patch images, highlighting considerable staining variations across KHP, NKI, BCI, EPFL and SGK cohorts. Columns two through five display the patches after normalisation using the Reinhard, Vahadane, Macenko, and StainGAN methods, demonstrating varying abilities to mitigate staining variations.
