## Supplementary Fig. 6 for "Normal Breast Tissue (NBT)-Classifiers: Advancing Compartment Classification in Normal Breast Histology"

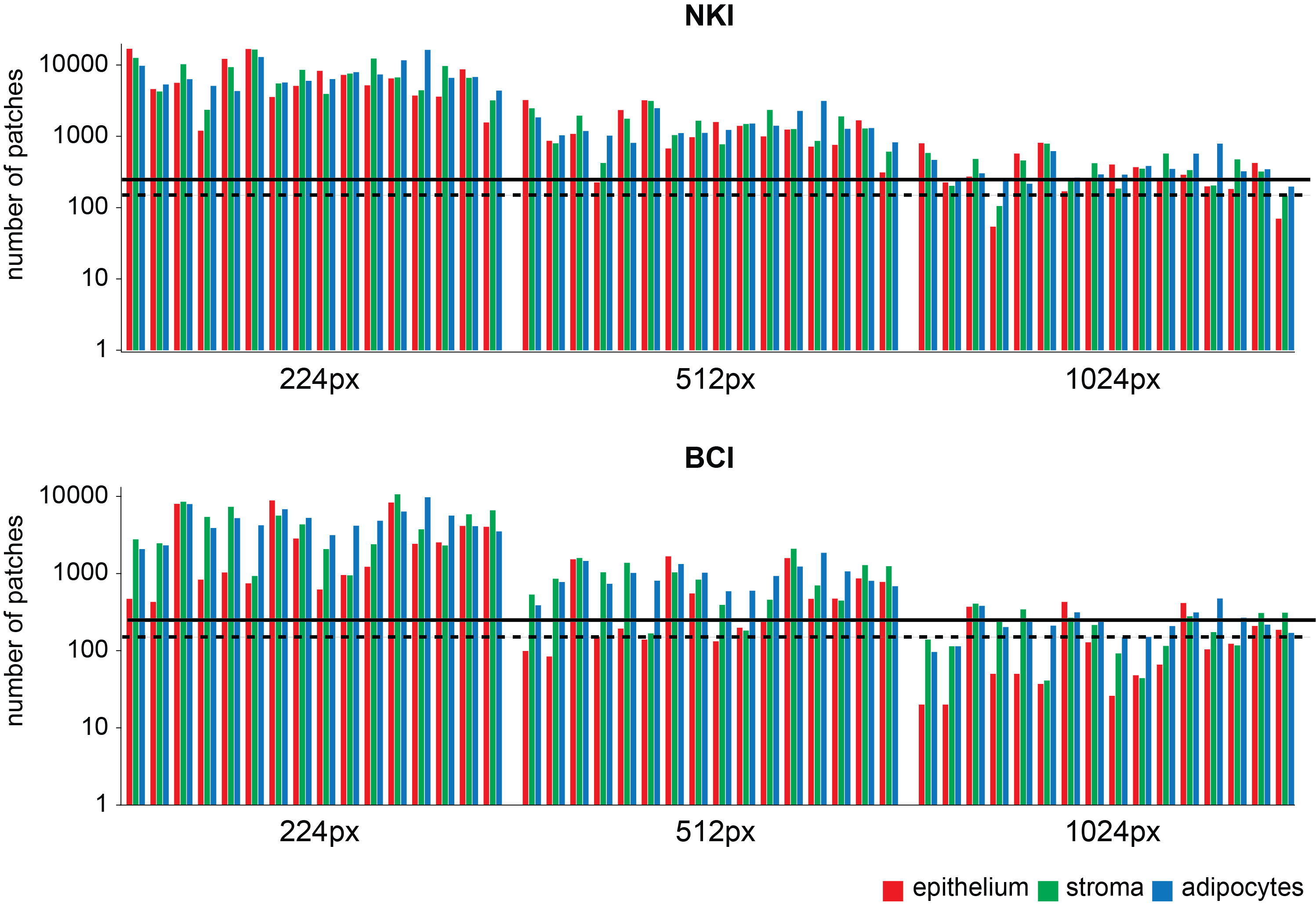

**Supplementary Fig. 6 Barplot showing the number of patches obtained using different patch sizes.** The panels illustrate the variation in the number of patches obtained from the NKI (top) and BCI (bottom) cohorts, using patch sizes of 224 x 224 pixels (denoted as 224px) (left), 512 x 512 pixels (denoted as 512px) (middle), and 1024 x 1024 pixels (denoted as 1024px) (right) at 40x magnification. In each panel, the x-axis represents individual WSI, with the coloured bar plots separately showing the number of epithelium (red), stroma (green), and adipocytes (blue) patches generated from each WSI at the corresponding patch size. The horizontal line indicates the number of 250 patches, while the horizontal dashed line indicates the number of 150 patches.
