## Supplementary Fig. 7 for "Normal Breast Tissue (NBT)-Classifiers: Advancing Compartment Classification in Normal Breast Histology"

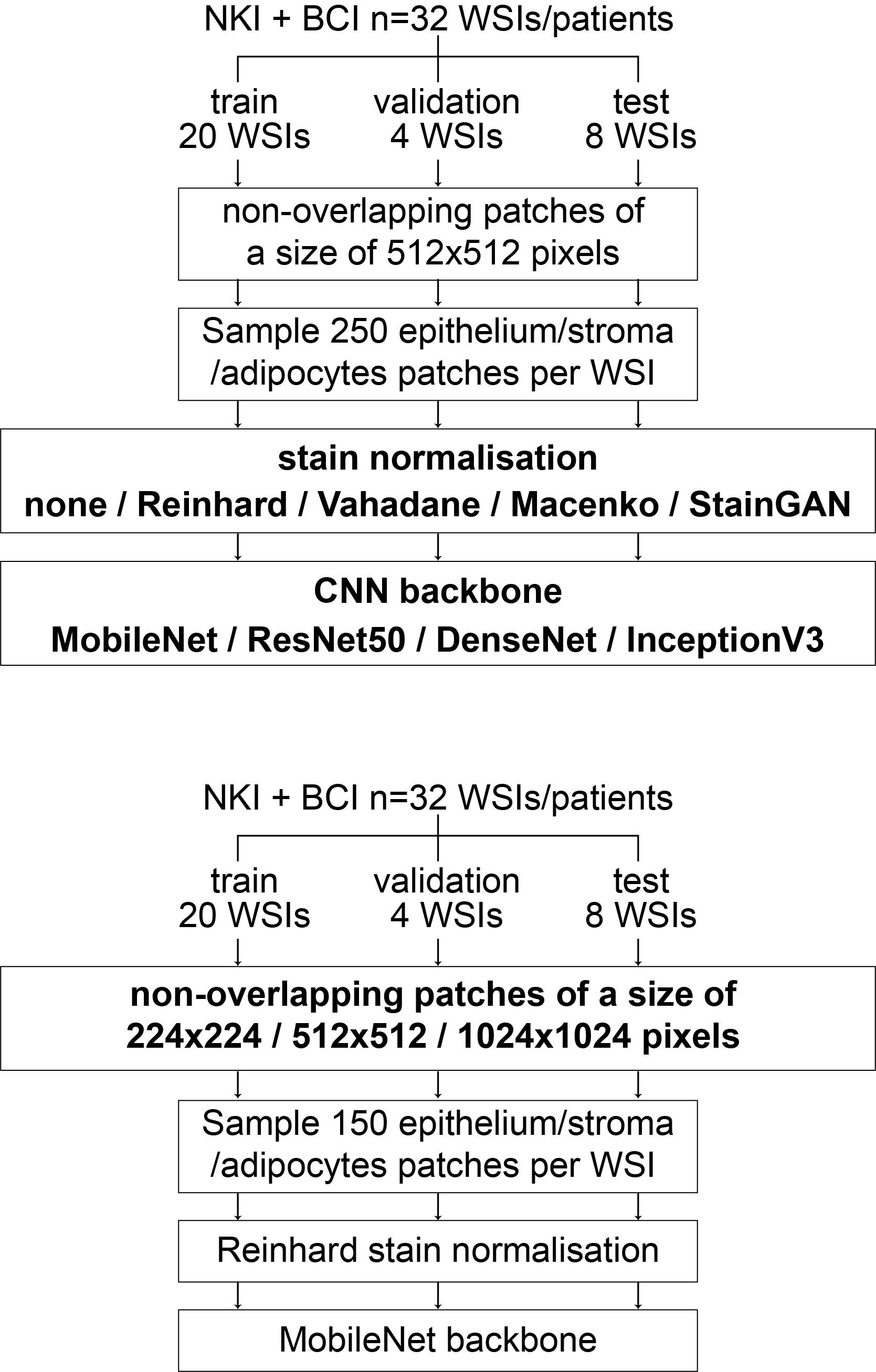

**Supplementary Fig. 7 Sample workflow for three-fold cross-validation experiments.** A total of 23 training configurations were evaluated, varying in stain normalisation methods, CNN backbones, and patch sizes.
