## Supplementary Fig. 8 for "Normal Breast Tissue (NBT)-Classifiers: Advancing Compartment Classification in Normal Breast Histology"

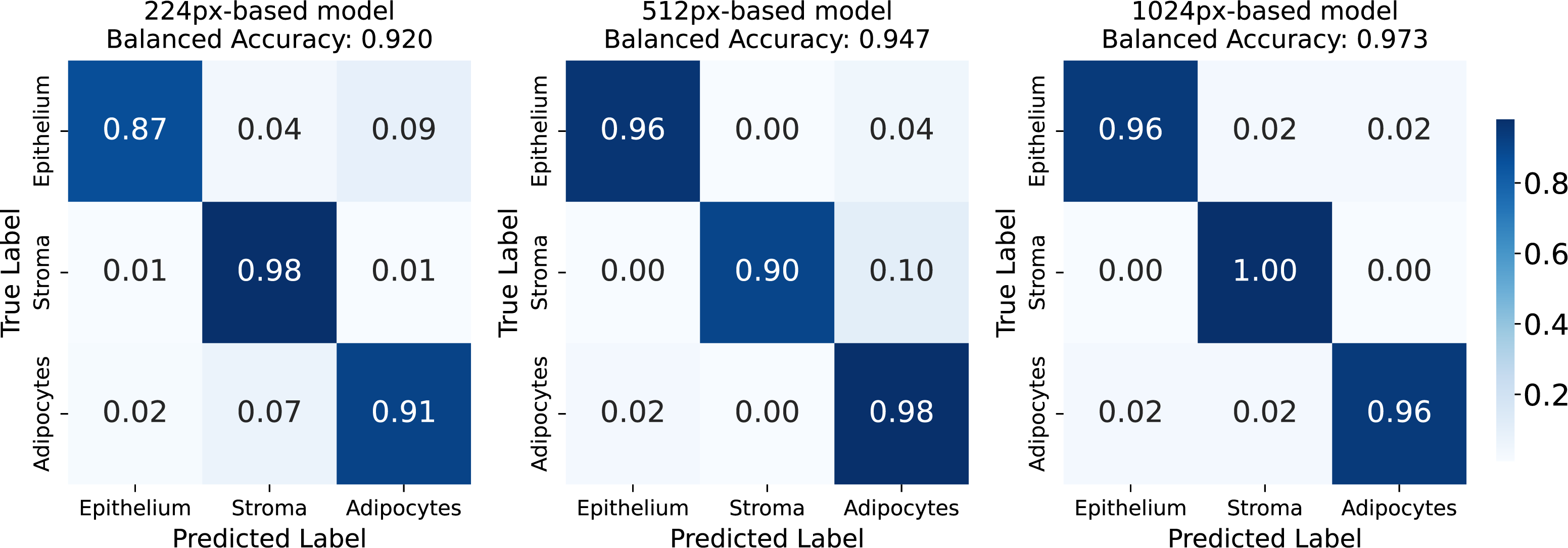

**Supplementary Fig. 8 Confusion matrices evaluating the 224px-, 512px and 1024px-based NBT-Classifiers on external datasets.** Each confusion matrix summarizes the classification results across the three external cohorts (KHP, EPFL, and SGK), comparing the predicted tissue classes: epithelium, stroma, and adipocytes (columns) against the ground-truth labels (rows). Diagonal cells indicate the number of correct predictions for each class, while off-diagonal cells indicate misclassifications. The colour intensity of each cell reflects the classification frequency, with lighter shades indicating higher counts. The overall balanced accuracy is derived from these matrixes and displayed at the top.
