## Supplementary Fig. 9 for "Normal Breast Tissue (NBT)-Classifiers: Advancing Compartment Classification in Normal Breast Histology"

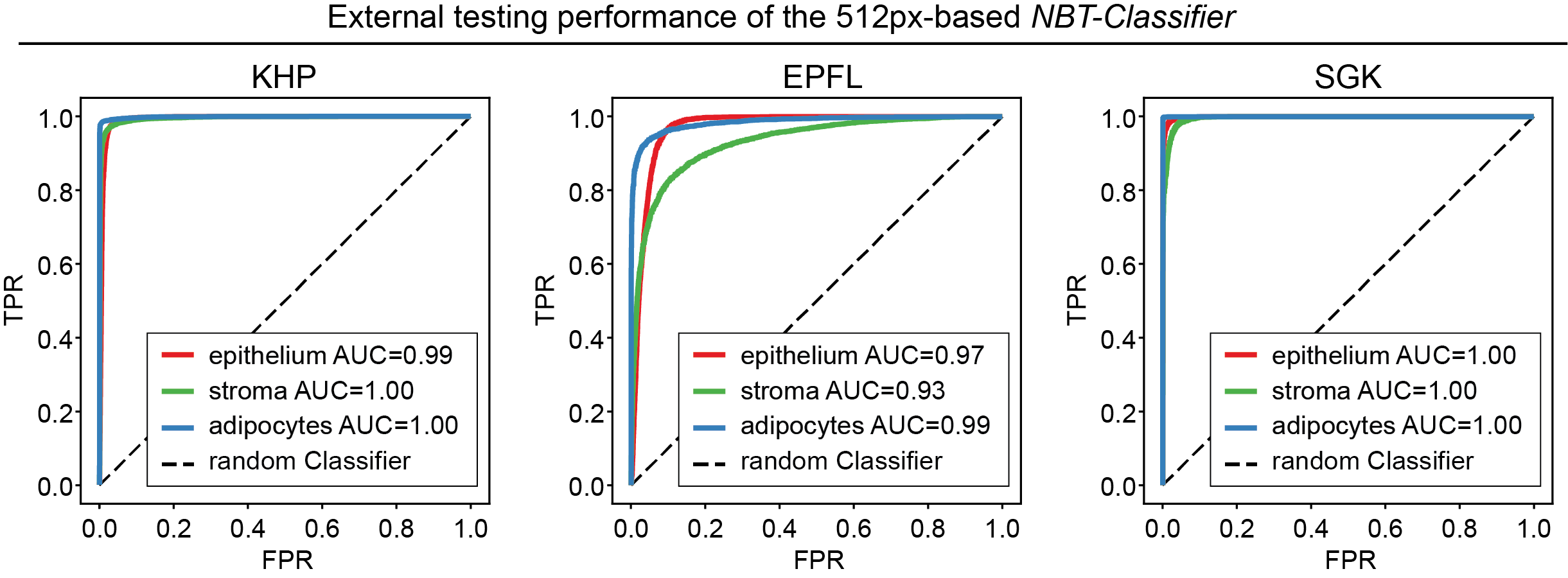

**Supplementary Fig. 9 Cohort-specific multi-class receiver operating characteristic curves for the 512px-based *NBT-Classifier*.** Panels from left to right show the receiver operating characteristic (ROC) curves of the 512px-based *NBT-Classifier* evaluated in the KHP, EPFL and SGK cohort, respectively. Performance of epithelium, stroma and adipocytes are indicated in red, green, and blue, respectively, with corresponding values of area under the ROC curve (AUC) presented below each curve.
