## Supplementary Fig. 10 for "Normal Breast Tissue (NBT)-Classifiers: Advancing Compartment Classification in Normal Breast Histology"

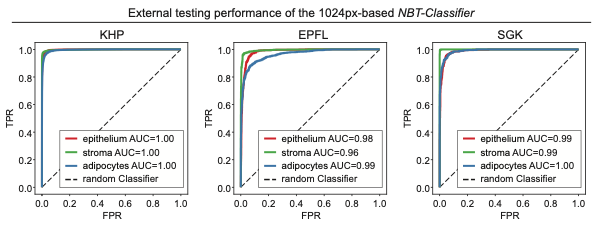

**Supplementary Fig. 10 Cohort-specific multi-class ROC curves for the 1024px-based *NBT-Classifier*.** Panels from left to right show the ROC curves of the 1024px-based *NBT-Classifier* evaluated in the KHP, EPFL and SGK cohort, respectively. Performance of epithelium, stroma and adipocytes are indicated in red, green, and blue, respectively, with corresponding values of AUC presented below each curve.
