## Supplementary Fig. 11 for "Normal Breast Tissue (NBT)-Classifiers: Advancing Compartment Classification in Normal Breast Histology"

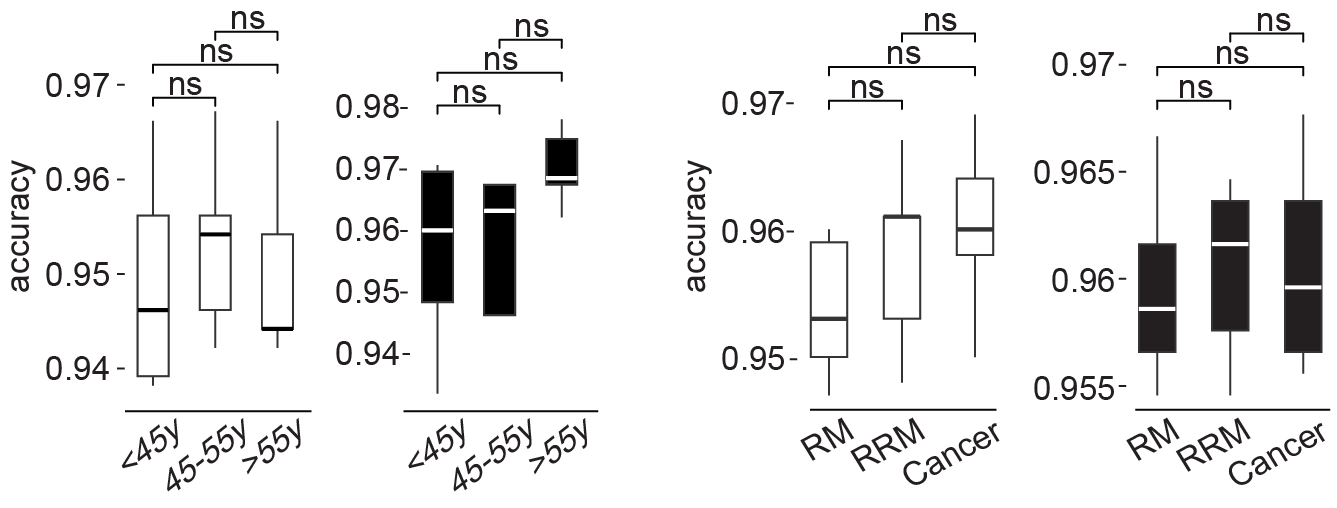

**Supplementary Fig. 11 Boxplot comparisons of *NBT-Classifiers’* performance across different age groups and NBT sources.** The performance of the 512px- and 1024px-based models is depicted in white and black boxplots, respectively. In each boxplot, the central line indicates the median, while the whiskers extend to the minimum and maximum values. The label "ns" denotes statistical non-significance, with adjusted p-values less than 0·05 considered significant. RM stands for reduction mammoplasty, and RRM stands for risk-reducing mastectomy.
