## Supplementary Fig. 12 for "Normal Breast Tissue (NBT)-Classifiers: Advancing Compartment Classification in Normal Breast Histology"

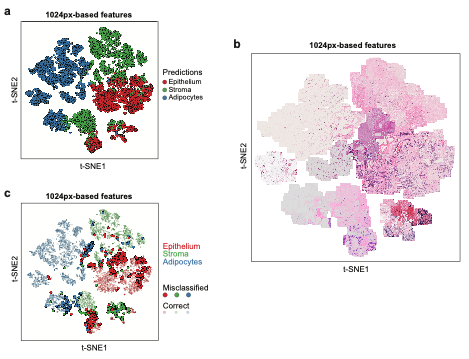

**Supplementary Fig. 12 t-SNE visualisation of features extracted by the 1024px-based *NBT-Classifier*.** Features are derived from the last layer (the layer before the output layer) of the 1024px-based *NBT-Classifier*. Panel **a** displays a t-Distributed Stochastic Neighbour Embedding (t-SNE) plot coloured by the 1024px-based *NBT-Classifier’s* predictions. Panel **b** overlays original H&E-stained patches onto a subset of random features (500 per tissue class) on the t-SNE plot. Panel **c** displays a t-SNE plot with points coloured according to ground-truth tissue classes. Misclassified patches are marked as opaque circles (alpha=1.0), while correctly classified patches are represented by semi-transparent circles (alpha=0.3).
