## Supplementary Fig. 13 for "Normal Breast Tissue (NBT)-Classifiers: Advancing Compartment Classification in Normal Breast Histology"

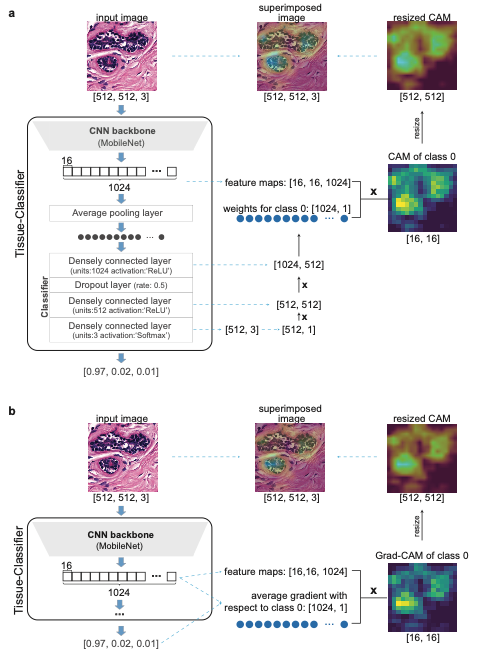

**Supplementary Fig. 13 Illustration of class activation mapping and gradient-weighted class activation mapping visualisations.** Class activation mapping (CAM) (**a**) uses class-specific weights to aggregate feature maps from the last convolutional layer of the CNN backbone, while gradient-weighted class activation mapping (Grad-CAM) (**b**) computes the average gradients. Both methods highlight high-attention regions that are relevant to a specific tissue class.
