## Supplementary Fig. 14 for "Normal Breast Tissue (NBT)-Classifiers: Advancing Compartment Classification in Normal Breast Histology"

**Supplementary Fig. 14 Additional CAM and Grad-CAM visualisations.** Panel **a** displays Grad-CAM heatmaps for 512 x 512-pixel patches (0·25 µm/pixel) of epithelium (top row), stroma (middle row), and adipocytes (bottom row). Panel **b** shows original 1024 x 1024-pixel H&E patches (0·25 µm/pixel) and predictions (top), CAM (bottom left) and Grad-CAM (bottom right) visualisations. Each row, from left to right, includes the original CAM or Grad-CAM heatmaps, resized heatmaps, and heatmaps overlaid on the original patches.
