## Supplementary Fig. 15 for "Normal Breast Tissue (NBT)-Classifiers: Advancing Compartment Classification in Normal Breast Histology"

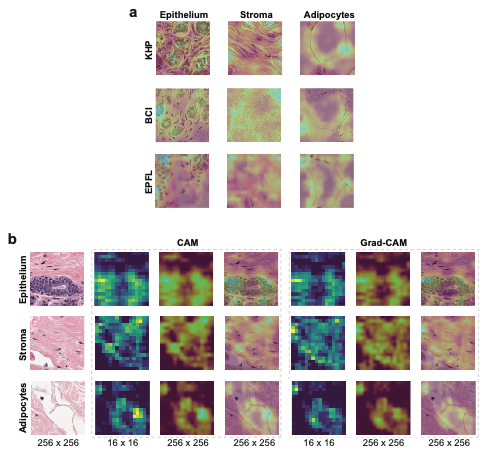

**Supplementary Fig. 15 Consistent histopathological patterns are captured across multiple cohorts.** In panel **a**, rows indicate CAM heatmaps overlaid on the Reinhard normalised epithelium, stroma and adipocytes patches (512 x 512 pixels, 0·25 µm/pixel) from KHP, BCI and EPFL cohorts. Each patch was predicted by the 512px-based *NBT-Classifier* with a probability of 1·0 of the predicted tissue class. The highlighted histopathological patterns in each tissue class are consistent across multiple cohorts. In panel **b**, patches are 256 x 256 pixels, 0·5 µm/pixel, obtained from WSIs in the SGK cohort that were scanned at 20x magnification, with CAM (left) and Grad-CAM (right) visualisations.
