## Supplementary Fig. 16 for "Normal Breast Tissue (NBT)-Classifiers: Advancing Compartment Classification in Normal Breast Histology"

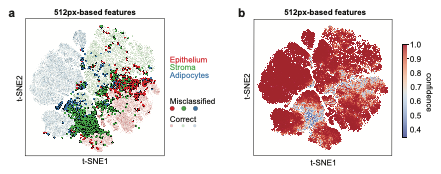

**Supplementary Fig. 16 t-SNE visualisation of features extracted by the 512px-based *NBT-Classifier*.** Features are derived from the last layer (just before the output layer) of the model. Panel **a** displays a t-SNE plot coloured by ground-truth tissue classes. Misclassified patches are marked as opaque circles (alpha=1.0), while correctly classified patches are represented by semi-transparent circles (alpha =0.3). Panel **b** presents the t-SNE visualisation of prediction confidence for the 512px-based *NBT-Classifier*. Each patch is coloured according to the probability of its predicted tissue class.
