## Supplementary Fig. 17 for "Normal Breast Tissue (NBT)-Classifiers: Advancing Compartment Classification in Normal Breast Histology"

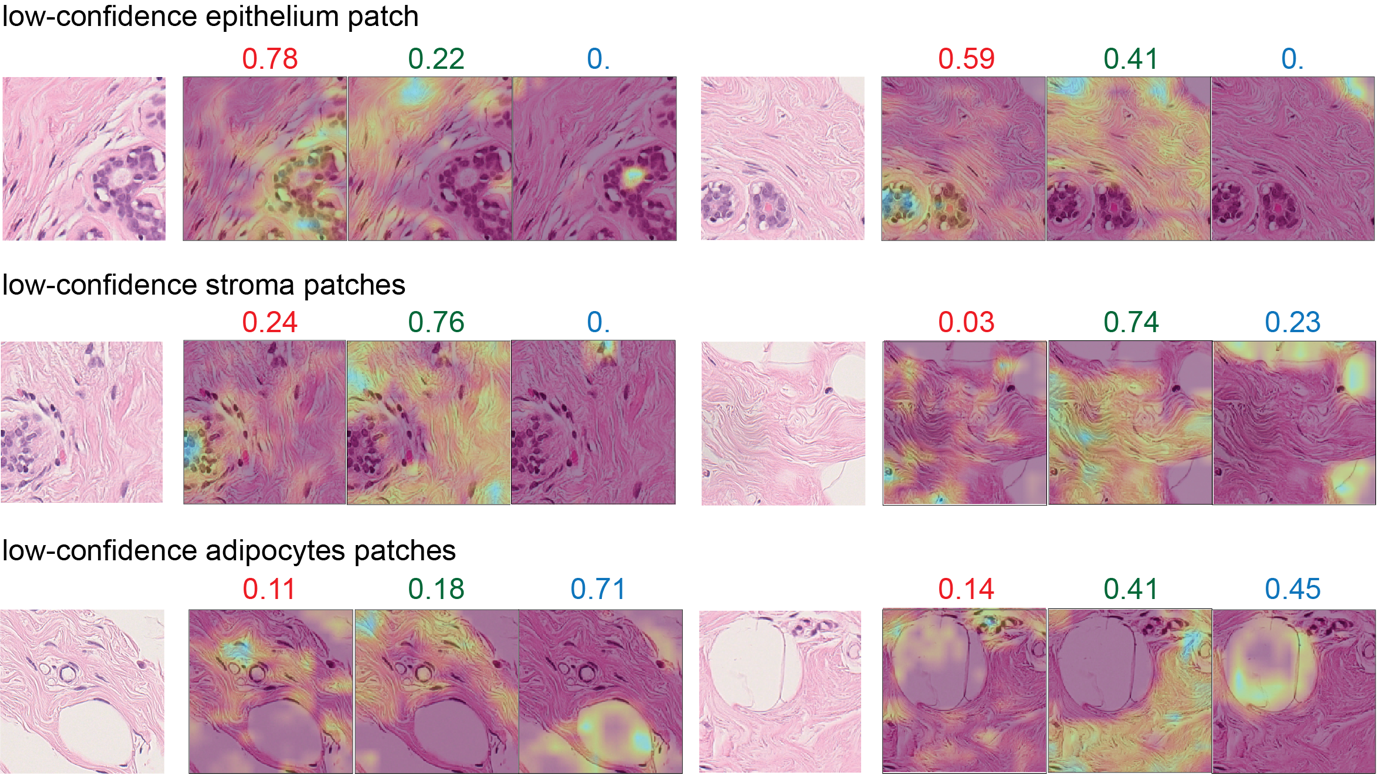

**Supplementary Fig. 17 CAM-based interpretation of low-confidence predictions by the 512px-based *NBT-Classifier*.** The rows, from top to bottom, show low-confidence 512 x 512-pixel patches predicted as epithelium, stroma, and adipocytes, respectively. For each patch, CAM heatmaps are generated for all three tissue classes and overlaid on the corresponding H&E patch image, highlighting class-specific high-attention regions. The predicted class probabilities are displayed at the top, with epithelium in red, stroma in green, and adipocytes in blue.
