## Supplementary Fig. 18 for "Normal Breast Tissue (NBT)-Classifiers: Advancing Compartment Classification in Normal Breast Histology"

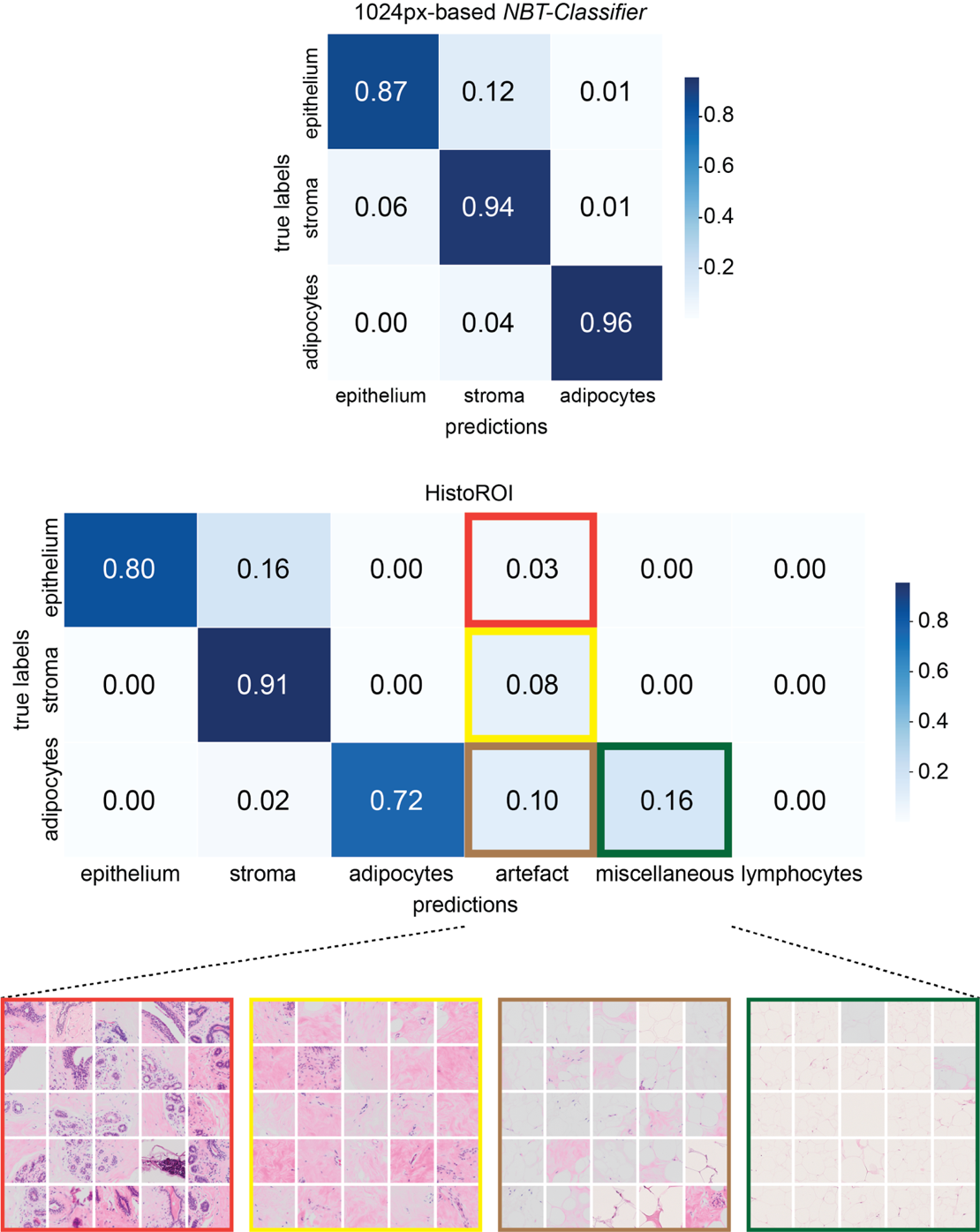

**Supplementary Fig. 18 Proportional confusion matrices for the 1024px-based *NBT-Classifier* and HistoROI.** For HistoROI, representative examples of epithelium and misclassified as artefact are indicated in red, stroma and misclassified as artefact are indicated in yellow, adipocytes and misclassified as artefact are indicated in brown, and adipocytes and misclassified as miscellaneous are indicated in green.
