## Supplementary Fig. 19 for "Normal Breast Tissue (NBT)-Classifiers: Advancing Compartment Classification in Normal Breast Histology"

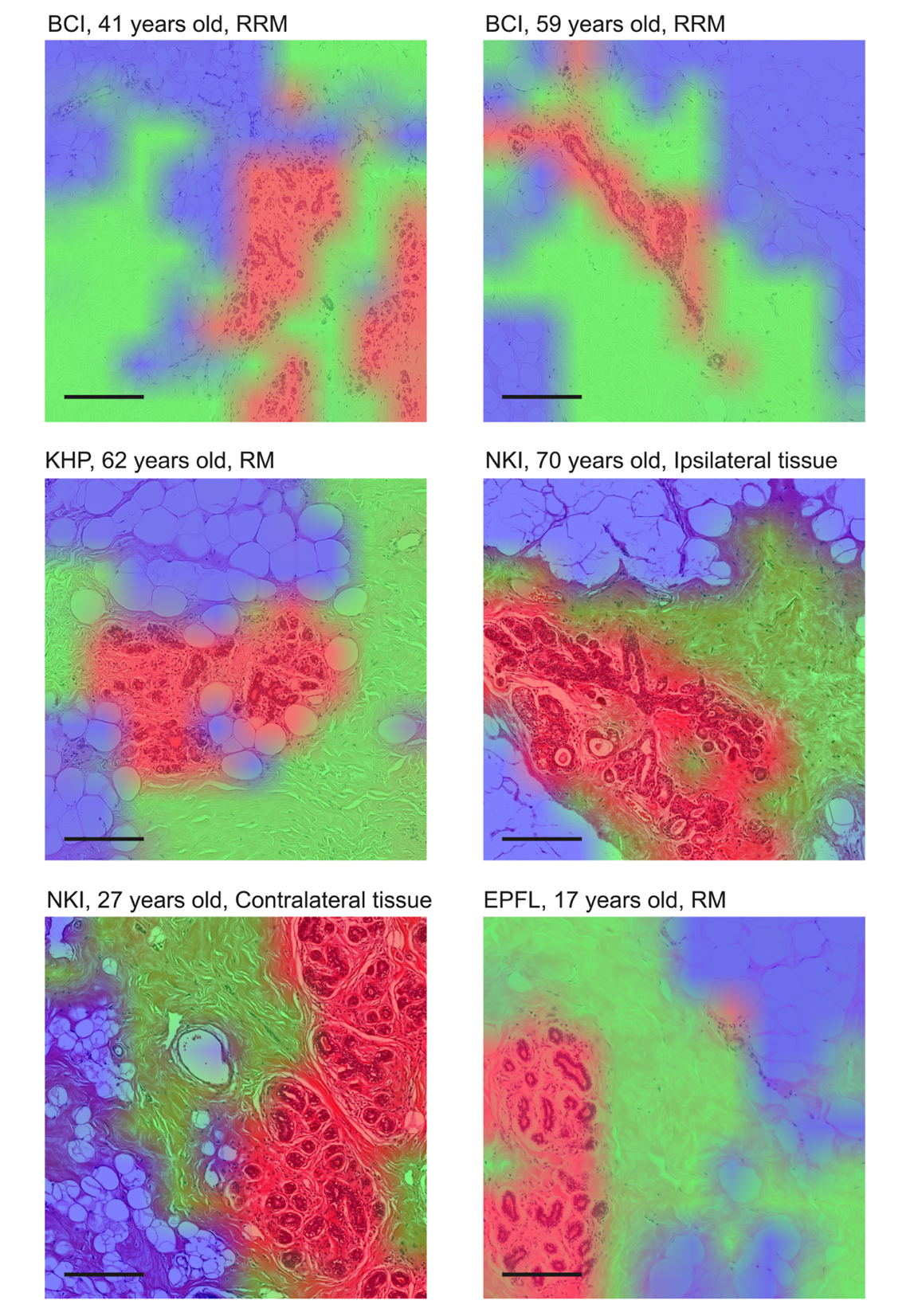

**Supplementary Fig. 19 Representative regions with overlaid tissue probability heatmaps.** Heatmaps were generated from NBTs with distinct patient ages and NBT sources, as indicated at the top of each figure. RRM: risk-reducing mastectomy; RM: reduction mammoplasty. Each heatmap clearly delineates the epithelium (red), stroma (green), and adipocytes (blue). Scale bar, 250 µm.
