## Supplementary Fig. 20 for "Normal Breast Tissue (NBT)-Classifiers: Advancing Compartment Classification in Normal Breast Histology"

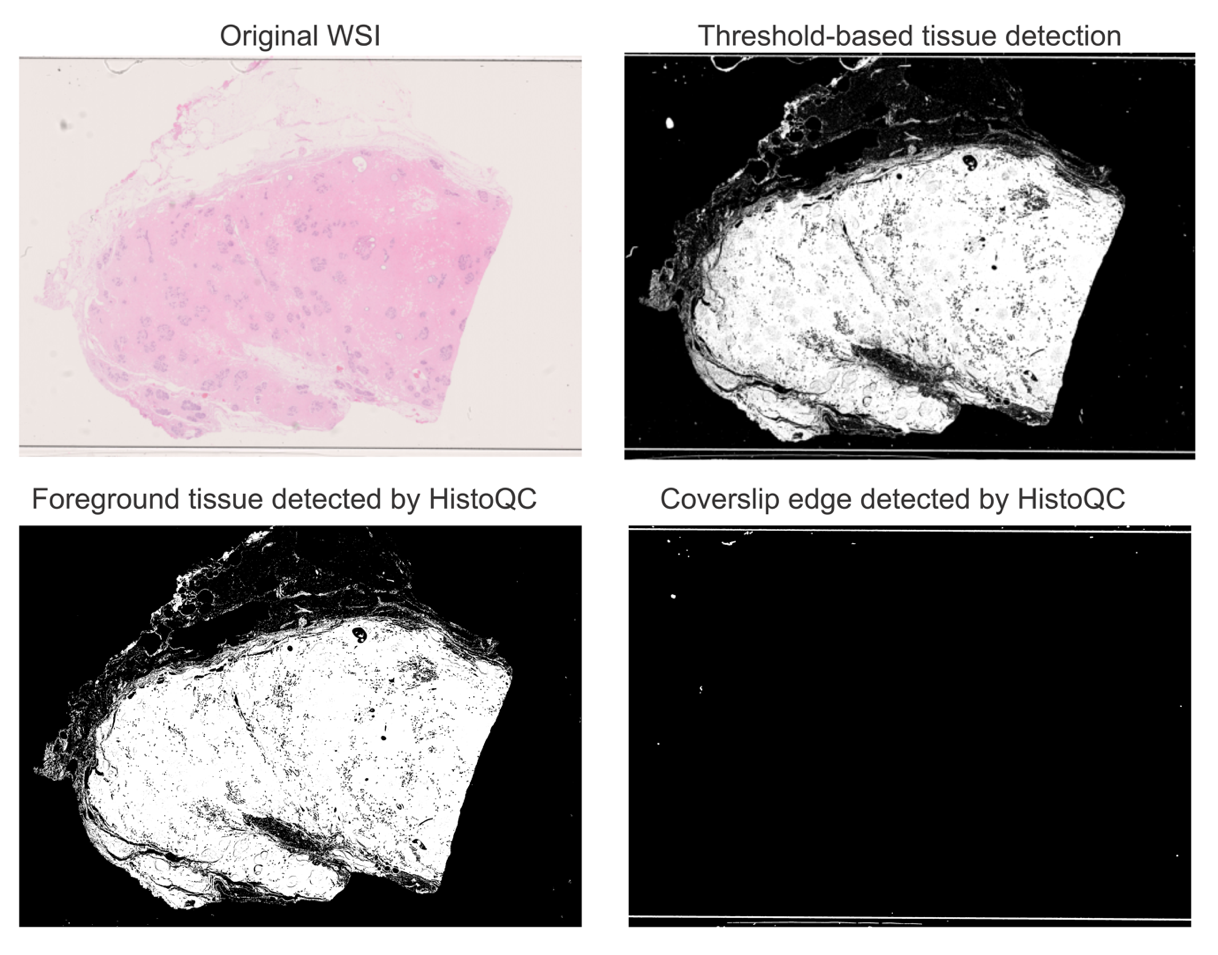

**Supplementary Fig. 20 Illustration of foreground tissue detection using HistoQC.** The top left image shows the original H&E-stained WSI of NBTs. The top right image shows the foreground tissue (in white) detected by the conventional Otsu threshold algorithm. Notably, the coverslip edge is incorrectly included as part of the foreground tissue. The bottom left image shows the foreground tissue (in white) detected by HistoQC, with the coverslip edge (in white) separately identified in the bottom right image.
