## Supplementary Fig. 21 for "Normal Breast Tissue (NBT)-Classifiers: Advancing Compartment Classification in Normal Breast Histology"

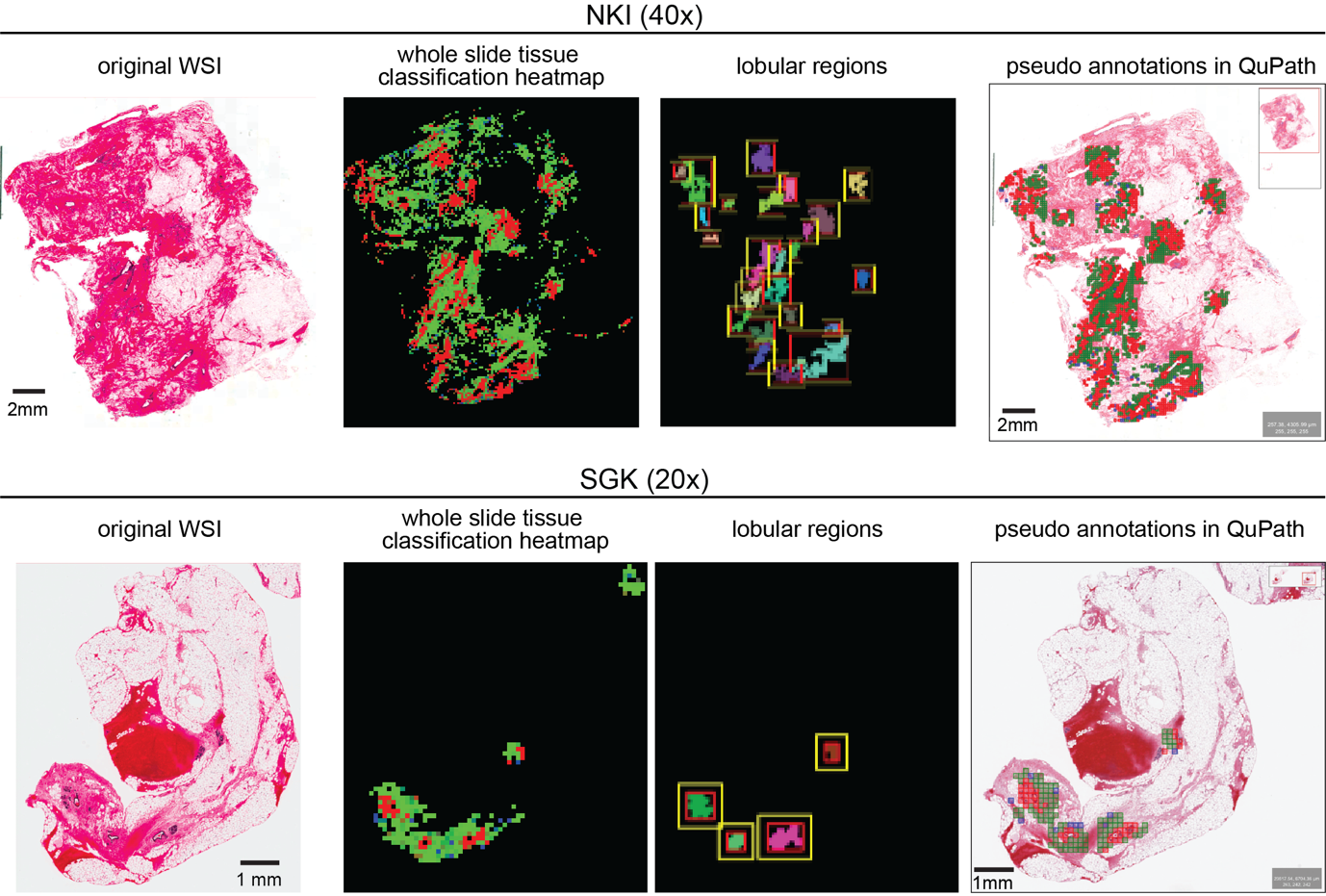

**Supplementary Fig. 21 Examples illustrating the proposed *NBT-Classifier*-based WSI pre-processing pipeline.** The top row shows a WSI example from the NKI cohort, scanned at 40x magnification, while the bottom row shows a WSI example from the SGK cohort, scanned at 20x magnification. From left to right, the images display the original WSI, the whole slide tissue classification heatmap, detected lobular regions (lobules are localised by the inner red boxes and peri-lobular regions are localised by the outer yellow boxes), and pseudo patch-level annotations (red for epithelium patches and green for stroma patches) visualised in QuPath v0.3.0.
