## Supplementary Fig. 22 for "Normal Breast Tissue (NBT)-Classifiers: Advancing Compartment Classification in Normal Breast Histology"

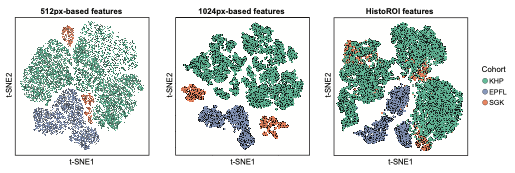

**Supplementary Fig. 22 t-SNE visualisation of feature embeddings illustrating domain shift across cohorts.** Feature embeddings are coloured by cohort for the 512px-based *NBT-Classifier* (left), 1024px-based *NBT-Classifier* (middle) and HistoROI (right).
