## Supplementary Table 1 for "Normal Breast Tissue (NBT)-Classifiers: Advancing Compartment Classification in Normal Breast Histology"

| **Supplementary Table 1. Overview of annotated WSI datasets of NBTs.** | | | | | |
| --- | --- | --- | --- | --- | --- |
| **WSI id** | **Magnification** | **Data split** | **Cohort** | **Type of specimen** | **Patient** |
|  |  |  |  |  | **age** |
| LS13-03088N dc | 40x | training | BCI | reduction mammoplasty | 24 |
| LS15-02541N-y |  |  |  |  | 40 |
| LS15-02672N-n |  |  |  |  | 40 |
| 3046 B2 |  |  |  |  | 55 |
| 2282 B6 |  |  |  | risk-reducing mastectomy | 29 |
| 1097 A7 |  |  |  |  | 32 |
| 1438 K2 |  |  |  |  | 34 |
| 1130 D6 |  |  |  |  | 36 |
| 3154 LEFT PM |  |  |  |  | 37 |
| 3239 Left PM |  |  |  |  | 38 |
| 3031PM A6 |  |  |  |  | 39 |
| 1991PM A2 |  |  |  |  | 41 |
| 2243 A2 |  |  |  |  | 41 |
| 1524 A3 |  |  |  |  | 45 |
| 2204 B8 |  |  |  |  | 46 |
| 2975PM A3 |  |  |  |  | 59 |
| T05-00863 A2 HE | 40x | training | NKI | Contralateral NBT | 27 |
| T10-09302 A2 HE |  |  |  |  | 32 |
| T12-04313 A1 HE |  |  |  |  | 36 |
| T10-10050 A3 HE |  |  |  |  | 40 |
| T04-10775 A5 HE |  |  |  |  | 47 |
| T10-10006 A3 HE |  |  |  |  | 56 |
| T16-00449 A1 HE |  |  |  | Ipsilateral NBT | 70 |
| T13-05327 A4 HE |  |  |  | risk-reducing mastectomy | 26 |
| T18-08826 I1 HE |  |  |  |  | 29 |
| T19-02002 I1 HE |  |  |  |  | 31 |
| T15-08578 A1 HE |  |  |  |  | 35 |
| T15-00483 A3 HE |  |  |  |  | 37 |
| T13-04640 A2 HE |  |  |  |  | 42 |
| T13-04670 A1 HE |  |  |  |  | 44 |
| T13-04602 A4 HE |  |  |  |  | 47 |
| T14-01017 A3 HE |  |  |  |  | 58 |
| Human_195_s01_HE | 40x | testing | EPFL | reduction mammoplasty | 16 |
| Human_103_s01_HE |  |  |  |  | 17 |
| Human_188_s01_HE |  |  |  |  | 17 |
| Human_129_s01_HE |  |  |  |  | 18 |
| Human_035_s01_HE |  |  |  |  | 26 |
| Human_016_s01_HE |  |  |  |  | 28 |
| Human_027_s01_HE |  |  |  |  | 31 |
| Human_172_s01_HE |  |  |  |  | 32 |
| Human_190_s01_HE |  |  |  |  | 33 |
| Human_131_s01_HE |  |  |  |  | 39 |
| 19001626_FPE_3 | 40x | testing | KHP | reduction mammoplasty | 22 |
| 18001177_FPE_1 |  |  |  |  | 23 |
| 17064108_FPE_1 |  |  |  |  | 29 |
| 18000951_FPE_2 |  |  |  |  | 30 |
| 17063396_FPE_5 |  |  |  |  | 32 |
| 17063838_FPE_2 |  |  |  |  | 37 |
| 17063839_FPE_3 |  |  |  |  | 37 |
| 17064113_FPE_4 |  |  |  |  | 38 |
| 19004666_FPE_9 |  |  |  |  | 41 |
| 17063968_FPE_3 |  |  |  |  | 45 |
| 17064240_FPE_4 |  |  |  |  | 47 |
| 17064241_FPE_7 |  |  |  |  | 47 |
| 17063451_FPE_3 |  |  |  |  | 59 |
| 17063504_FPE_6 |  |  |  |  | 62 |
| 17063106_FPE_3 |  |  |  |  | 69 |
| 17063107_FPE_4 |  |  |  |  | 69 |
| K104210 | 20x | testing | SGK | core biopsies from healthy donors | 19 |
| K106560 |  |  |  |  | 29 |
| K102595 |  |  |  |  | 30 |
| K106776 |  |  |  |  | 32 |
| K102518 |  |  |  |  | 40 |
| K108158 |  |  |  |  | 46 |
| K104934 |  |  |  |  | 50 |
| K108301 |  |  |  |  | 51 |
| K102425 |  |  |  |  | 52 |
| K104724 |  |  |  |  | 60 |
| K108355 |  |  |  |  | 72 |
| K106822 |  |  |  |  | 74 |

Note: BCI: the Barts Cancer Institute in London (UK); NKI: the Netherlands Cancer Institute in Amsterdam (Netherlands); EPFL: the École Polytechnique Fédérale de Lausanne in Lausanne (Switzerland); KHP: the King's Health Partners Cancer Biobank in London (UK); SGK: the publicly available Susan G. Komen Tissue Bank; NBT: normal breast tissue.
