## Supplementary Table 2 for "Normal Breast Tissue (NBT)-Classifiers: Advancing Compartment Classification in Normal Breast Histology"

### **Supplementary Table 2. Summary of annotated patch-level datasets.**

| **Patch-level**  **dataset** | **Data split** | **Cohort** | **Patch size** | **Annotation** | **Number of patches** |
| --- | --- | --- | --- | --- | --- |
| NKI_512 | training | NKI | 512 | epithelium | 21,241 |
| NKI_512 | training | NKI | 512 | stroma | 23,735 |
| NKI_512 | training | NKI | 512 | adipocytes | 23,516 |
| NKI_224 | training | NKI | 224 | epithelium | 110,639 |
| NKI_224 | training | NKI | 224 | stroma | 123,497 |
| NKI_224 | training | NKI | 224 | adipocytes | 122,668 |
| NKI_1024 | training | NKI | 1024 | epithelium | 5,322 |
| NKI_1024 | training | NKI | 1024 | stroma | 5,881 |
| NKI_1024 | training | NKI | 1024 | adipocytes | 5,900 |
| BCI_512 | training | BCI | 512 | epithelium | 9,144 |
| BCI_512 | training | BCI | 512 | stroma | 14,214 |
| BCI_512 | training | BCI | 512 | adipocytes | 15,273 |
| BCI_224 | training | BCI | 224 | epithelium | 47,397 |
| BCI_224 | training | BCI | 224 | stroma | 71,889 |
| BCI_224 | training | BCI | 224 | adipocytes | 79,138 |
| BCI_1024 | training | BCI | 1024 | epithelium | 2,281 |
| BCI_1024 | training | BCI | 1024 | stroma | 3,212 |
| BCI_1024 | training | BCI | 1024 | adipocytes | 3,768 |
| KHP_512 | external validation | KHP | 512 | epithelium | 16,995 |
| KHP_512 | external validation | KHP | 512 | stroma | 14,430 |
| KHP_512 | external validation | KHP | 512 | adipocytes | 20,021 |
| KHP_1024 | external validation | KHP | 1024 | epithelium | 4,039 |
| KHP_1024 | external validation | KHP | 1024 | stroma | 3,327 |
| KHP_1024 | external validation | KHP | 1024 | adipocytes | 4,467 |
| EPFL_512 | external validation | EPFL | 512 | epithelium | 2,813 |
| EPFL_512 | external validation | EPFL | 512 | stroma | 5,378 |
| EPFL_512 | external validation | EPFL | 512 | adipocytes | 4,947 |
| EPFL_1024 | external validation | EPFL | 1024 | epithelium | 654 |
| EPFL_1024 | external validation | EPFL | 1024 | stroma | 1,149 |
| EPFL_1024 | external validation | EPFL | 1024 | adipocytes | 1,068 |
| SGK_256 | external validation | SGK | 256 | epithelium | 1,674 |
| SGK_256 | external validation | SGK | 256 | stroma | 1,908 |
| SGK_256 | external validation | SGK | 256 | adipocytes | 3,034 |
| SGK_512 | external validation | SGK | 512 | epithelium | 416 |
| SGK_512 | external validation | SGK | 512 | stroma | 472 |
| SGK_512 | external validation | SGK | 512 | adipocytes | 741 |

Note: “train” refers to three-fold cross-validation, while “test” refers to external validation. The patch resolution is 0·5 µm per pixel for the SGK cohort and 0·25 µm per pixel for other cohorts. Patches were extracted without overlapping.
