## Supplementary Table 3 for "Normal Breast Tissue (NBT)-Classifiers: Advancing Compartment Classification in Normal Breast Histology"

### **Supplementary Table 3. Summary of three-fold cross validation accuracies.**

| **Cohort** | **Stain normalisation** | **CNN backbone** | **Patch size** | **Accuracy (mean)** |
| --- | --- | --- | --- | --- |
| NKI+BCI | none | MobileNet | 512 | 95.75% |
| NKI+BCI | Macenko | MobileNet | 512 | 97.32% |
| NKI+BCI | Vahadane | MobileNet | 512 | 97.05% |
| NKI+BCI | Reinhard | MobileNet | 512 | 97.72% |
| NKI+BCI | GAN | MobileNet | 512 | 96.96% |
| NKI+BCI | none | ResNet50 | 512 | 96.86% |
| NKI+BCI | Macenko | ResNet50 | 512 | 96.84% |
| NKI+BCI | Vahadane | ResNet50 | 512 | 96.77% |
| NKI+BCI | Reinhard | ResNet50 | 512 | 97.25% |
| NKI+BCI | GAN | ResNet50 | 512 | 96.98% |
| NKI+BCI | none | InceptionV3 | 512 | 95.90% |
| NKI+BCI | Macenko | InceptionV3 | 512 | 96.25% |
| NKI+BCI | Vahadane | InceptionV3 | 512 | 96.23% |
| NKI+BCI | Reinhard | InceptionV3 | 512 | 96.32% |
| NKI+BCI | GAN | InceptionV3 | 512 | 96.86% |
| NKI+BCI | none | DenseNet | 512 | 96.40% |
| NKI+BCI | Macenko | DenseNet | 512 | 97.08% |
| NKI+BCI | Vahadane | DenseNet | 512 | 96.49% |
| NKI+BCI | Reinhard | DenseNet | 512 | 96.79% |
| NKI+BCI | GAN | DenseNet | 512 | 96.58% |
| NKI+BCI | Reinhard | MobileNet | 224 | 92.63% |
| NKI+BCI | Reinhard | MobileNet | 512 | 95.53% |
| NKI+BCI | Reinhard | MobileNet | 1024 | 97.03% |
