## Supplementary Table 4 for "Normal Breast Tissue (NBT)-Classifiers: Advancing Compartment Classification in Normal Breast Histology"

### **Supplementary Table 4. A summary of studies of NBTs with manual annotations in recent ten years.**

| **Title** | **Source** | **Sample size** | **Manual annotations** | **Model** |
| --- | --- | --- | --- | --- |
| Deep learning assessment of breast terminal duct lobular unit involution: Towards automated prediction of breast cancer risk ([10.1371/journal.pone.0231653](https://doi.org/10.1371/journal.pone.0231653)) | benign breast  biopsies | 92 WSIs | Terminal duct lobular units (92 WSIs), acini and adipose tissue (50 WSIs) | Convolutional  neural network |
| Automated quantification of levels of breast terminal duct lobular (TDLU) involution using deep learning ([10.1038/s41523-021-00378-7](https://emckclac-my.sharepoint.com/personal/k21066795_kcl_ac_uk/Documents/PHD_Thesis/Thesis_manuscripts/NBT-Classifier/MCP_DigitalHealth/10.1038/s41523-021-00378-7)) | benign breast  biopsies | 33 WSIs | Epithelium (and myoepithelium),  extralobular stroma, intralobular  stroma, adipose tissue, lumens of acini, and small calibre blood vessels (“capillaries”) | Convolutional  neural network |
| Detection of lobular structures in normal breast tissue ([10.1016/j.compbiomed.2016.05.004](https://doi.org/10.1016/j.compbiomed.2016.05.004)) | Reduction  mammoplasty | 9 WSIs | Lobules and ducts | Mixed methods |
